# *IsoBayes*: a Bayesian approach for single-isoform proteomics inference

**DOI:** 10.1101/2024.06.10.598223

**Authors:** Jordy Bollon, Michael R Shortreed, Ben T Jordan, Rachel Miller, Erin Jeffery, Andrea Cavalli, Lloyd M Smith, Colin Dewey, Gloria M Sheynkman, Simone Tiberi

## Abstract

**Motivation:** Studying protein isoforms is an essential step in biomedical research; at present, the main approach for analyzing proteins is via bottom-up mass spectrometry proteomics, which return peptide identifications, that are indirectly used to infer the presence of protein isoforms. However, the detection and quantification processes are noisy; in particular, peptides may be erroneously detected, and most peptides, known as shared peptides, are associated to multiple protein isoforms. As a consequence, studying individual protein isoforms is challenging, and inferred protein results are often abstracted to the gene-level or to groups of protein isoforms.

**Results:** Here, we introduce *IsoBayes*, a novel statistical method to perform inference at the isoform level. Our method enhances the information available, by integrating mass spectrometry proteomics and transcriptomics data in a Bayesian probabilistic framework. To account for the uncertainty in the measurement process, we propose a two-layer latent variable approach: first, we sample if a peptide has been correctly detected (or, alternatively filter peptides); second, we allocate the abundance of such selected peptides across the protein(s) they are compatible with. This enables us, starting from peptide-level data, to recover protein-level data; in particular, we: i) infer the presence/absence of each protein isoform (via a posterior probability), ii) estimate its abundance (and credible interval), and iii) target isoforms where transcript and protein relative abundances significantly differ.

We benchmarked our approach in simulations, and in two multi-protease real datasets: our method displays good sensitivity and specificity when detecting protein isoforms, its estimated abundances highly correlate with the ground truth, and can detect changes between protein and transcript relative abundances.

**Availability and implementation:** *IsoBayes* is freely distributed as a Bioconductor R package, and is accompanied by an example usage vignette.

## 1 Introduction

Post-transcriptional regulatory mechanisms, such as alternative splicing or alternative promoter usage, allow single genes to code for multiple isoforms. For example, in humans, it was estimated that approximately 20,000 genes may give rise to over 300,000 protein isoforms (Deveson et al., 2018; Willyard, 2018). Characterization of protein isoform diversity, which includes the identification of physiologically relevant protein isoforms, as well as disease-associated aberrant splicing, is a crucial step in biomedical research. At present, the main strategy to infer proteins is via bottom-up mass spectrometry (MS) proteomics, where proteins are indirectly measured via peptides, which act as surrogate markers for their protein(s) of origin. Due to the high degree of sequence sim-ilarity between isoform sequences, most peptides, called shared peptides, are compatible with multiple protein isoforms; furthermore, the identification of peptides is a noisy process which can result in erroneous detections (Vesvizhskii and Aebersold, 2005). Therefore, inference at the isoform level is challenging, and results are often abstracted to the gene level, or to groups of multiple protein isoforms.

A few methods have been developed to perform inference at the isoform level; notably: *ProteinProphet* (Nesvizhskii et al., 2003), one of the first probabilistic methods for protein identification, that infers protein isoforms via an expectation-maximization algorithm based on a bipartite graph of peptides and associated proteins; *Fido* (Serang et al., 2010), *a Bayesian network model, that, via graph-transforming algorithms, groups and scores proteins based on peptide-spectrum matches; PIA* (Uszkoreit et al., 2015), which ranks protein isoforms based on the scores of the peptides they are compatible with; and *EPIFANY* (Pfeuffer et al., 2020), *that models the conditional dependencies between proteins and peptides via a Bayesian network representation, and estimates protein isoform posterior probabilities exploiting a loopy belief propagation algorithm combined with convolution trees. However, due to the prevalence of shared peptides, protein identification is affected by low statistical power; furthermore, inference only focuses on identifying protein isoforms (presence vs. absence), and not on further measures such as their abundance. In order to obtain more stable results, HIquant* (Bryan et al., 2016) infers ratios of protein-isoform abundances. However, *HIquant* does not estimate the presence/absence or the abundance of individual isoforms, and requires multiple samples (i.e., biological replicates); because of these limitations, it is therefore not applicable in the context of this work, which focuses on the inference of protein isoform presence and abundance, from individual MS samples.

Since transcriptomics protocols have better isoform-level resolution, compared to MS protocols, in recent years, some approaches have been proposed to enhance MS data with mRNA expression levels (Ramakrishnan et al., 2009; Ma et al., 2017, 2019; Liu et al., 2017; Carlyle et al., 2018; Salovska et al., 2020). This approach is motivated by the fact that mRNA is a prerequisite of protein, and their abundances are in general positively correlated (Lu et al., 2007; Maier et al., 2009; de Sousa Abreu et al., 2009; Edfors et al., 2016; Liu et al., 2016; Wang et al., 2019). Among these methods, Miller et al. (2022) used long-read RNA-seq data to enhance the reference isoform database, and recover protein groups with high mRNA abundance. Nonetheless, none of these frameworks achieves isoform resolution. Given the critical role of splicing in biology, more accurate methods for the detection of protein isoforms would be highly beneficial for life scientists.

Here, we present *IsoBayes*, a novel Bayesian approach to study protein isoforms from MS data, which also integrates transcriptomics data, when available.

## 2 Matherials and Methods

### 2.1 *IsoBayes* overview

Isoform-level data is characterized by two major sources of variability: biological noise, which is of interest, and technical noise, that is nuisance and is due to the measurement process. Our goal is to disentangle the two sources of uncertainty, in order to perform inference on the biological process; to this aim, we explicitly model the noise arising from both shared and erroneously detected peptides. In particular, isoform level abundance (and its presence/absence) is treated as a latent variable (i.e., an unknown parameter), and is sampled via a latent variable model.

Before analyzing the data, peptides are usually filtered based on a false discovery rate (FDR) threshold (typically 0.01); while this allows the removal of unreliable peptides, it also represents a crude cutoff. Here, we propose two distinct frameworks: one based on classical FDR filtering (that we call FDR mode), and a more complex one that uses the probability that peptides are correctly detected (called PEP mode). The FDR, peptide-error probability (PEP) and abundance of each peptide are taken as inputs here, and can be estimated by various proteomics tools, such as *MetaMorpheus* (Solntsev et al., 2018), *Percolator* (Käll et al., 2007; The et al., 2016) or *MaxQuant* (Cox and Mann, 2008). In our view, the PEP mode is a more accurate way of modelling peptide uncertainty, because it allows weighting peptides based on their probability of being correctly identified; furthermore, since more peptides are analyzed, inferential results are provided for a larger number of protein isoforms. Our latent variable approach works in 2 steps. In the first step, we either filter peptides based on their FDR (FDR mode), or sample if they are erroneously identified based on their error probability (PEP mode). In the second step, only for correct detections, we allocate the abundance of each peptide across the protein isoform(s) it is compatible with. This procedure, starting from peptide-level information, allows us to recover protein isoform-level abundance and presence.

When available, *IsoBayes* also allows for the integration of transcriptomics data. In particular, the relative abundance of transcript isoforms, estimated from (short or long-read) RNA-sequencing (RNA-seq) data, is used to formulate an informative prior for the relative abundance of the corresponding protein isoforms. Therefore, given a peptide associated to two isoforms, with high and low mRNA abundance, *a priori*, we assume that the peptide abundance is primarily coming from the first case. Clearly, this assumes a positive mRNA-protein correlation, with greater correlation leading to higher benefits. Overall, this integration enhances the data available to the model, and hence improves the accuracy of the inferential results.

For each protein isoform, our approach estimates both its presence and abundance, and provides a measure of the uncertainty of both estimates, via the posterior probability of presence, and a credible interval of its abundance. Abundance estimates, and respective credible intervals, are also aggregated at the gene-level. Additionally, when RNA-seq data is available, we study changes between isoform mRNA and protein relative abundances; in particular, for each isoform, we compute the log2-fold change (log2-FC) between protein and mRNA relative abundances, and estimate the probability that the relative abundance is higher at the protein-level than at the transcript-level. This feature allows scientists to identify candidate isoforms where protein and mRNA abundance levels may differ.

Furthermore, our tool is flexible and general: it requires peptide-level information (i.e., FDR, PSM counts or intensities, and optionally PEP), which can be obtained from both data dependent acquisition and data independent acquisition approaches, and from both label-free and labeled pipelines. Our method is also compatible with the output from any proteomics pipeline (e.g., *MetaMorpheus, Percolator*, and *MaxQuant* ) and, as a measure of abundance, users could employ either peptide intensities, or peptide spectral match (PSM) counts. Note that intensities or PSM counts are only proxies for the actual abundance of peptides, which is not exactly measured in MS data. In this manuscript, we use the term “abundance” to refer to those noisy measurements.

### 2.2 Mathematical modelling

Initially, we perform a trivial filtering step, where we remove protein isoforms which are not associated to any detected peptide, because they cannot be identified in the given dataset. All the inference is performed on protein isoforms which could potentially be detected, i.e. those associated to at least one (unique or shared) detected peptide, and results refer to them.

Given *P* such protein isoforms, we assume that the overall protein abundance, denoted by *n* ∈ N, is distributed across the *P* isoforms according to a multinomial distribution:

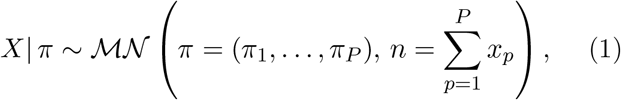

where *X* = (*X*_1_, … , *X*_*P*_ ), with *X*_*p*_ representing the random variable indicating the overall abundance originating from the *p*-th protein (and *x*_*p*_ its realization), and *π*_*p*_ is the probability that a unit of abundance comes from the *p*-th protein, with 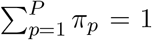. Our method is designed to work with integers: this is a convenience choice that simplifies the inference, particularly in the latent variable sampling. However, protein abundance in *X* can refer to either PSM counts, which are already discrete, or intensities, that are continuous. In the latter case, intensities are rounded to the closest integer, which introduces a minimal approximation. In our benchmarks (Section 3.5), we show that rounded intensities and PSM counts lead to similar inferential results.

If *X* was observed, the likelihood of the model could be easily computed as the density of the multinomial distribution in (1); however, measurements refer to peptides, and *X* is treated as a latent state. A graphical model of our method is displayed in Figure 1, and Supplementary Details report how the abundances of the *N* peptides, *Y* = (*Y*_1_, … , *Y*_*N*_ ), is obtained from the abundances of the *P* proteins, *X* = (*X*_1_, … , *X*_*P*_ ). Therefore, the likelihood is defined with respect to the actual observations (i.e., the peptide measurements), and can be written as an integral over the latent data: *L*(*π* |*Y* ) = ∫_*X*_ *f* (*Y, X* = *x*|*π*)*dx*, where *f* (*Y, X* = *x*|*π*) is the joint density of observations *Y* and latent states *X*, given parameters *π*. Here, instead of working with this integral, we employ a Bayesian data augmentation approach (Tanner and Wong, 1987; Gelfand and Smith, 1990a) where parameters and latent states are alternately sampled from their conditional distributions (see Section 2.6).

**Figure 1:**
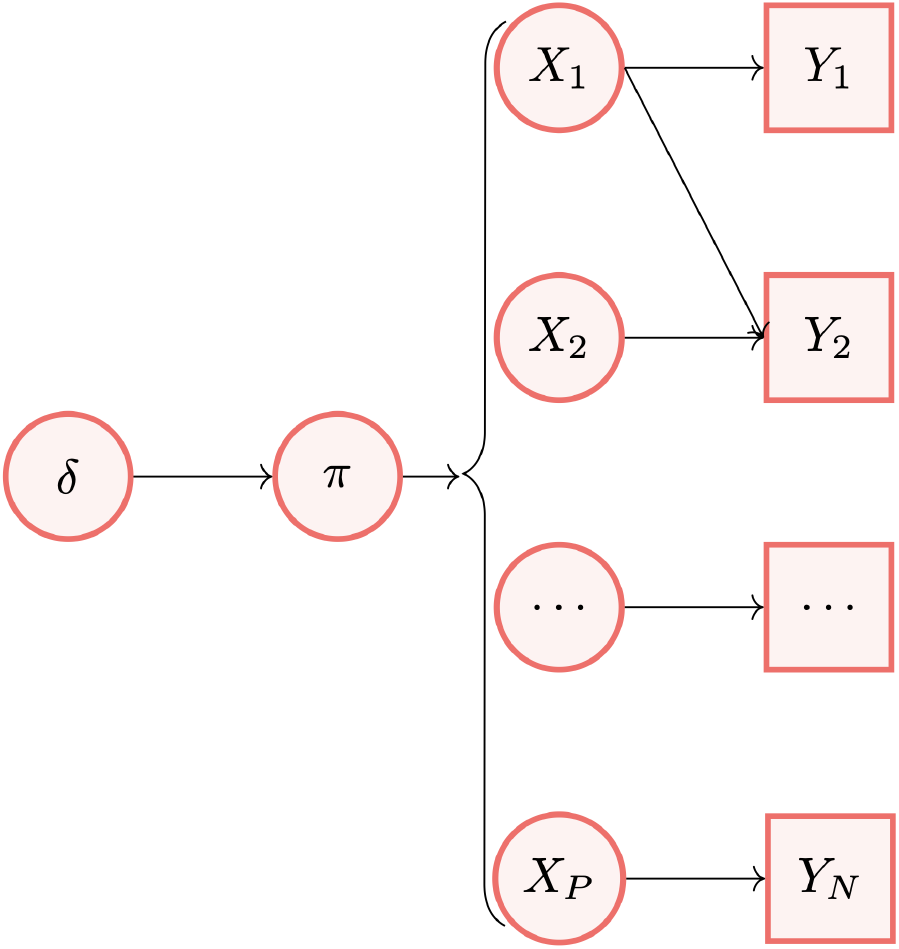
*IsoBayes* graphical model, connecting prior parameters *δ* to parameters *π*, then to latent protein abundances *X*, and finally to peptide-level observations *Y* . In the example above, protein 1 is connected to unique peptide 1 and to shared peptide 2, protein 2 is associated to shared peptite 2, while protein *P* is connected to unique peptide *N* . *IsoBayes*, starting from peptide-level data *Y* , aims to recover the unobserved protein-level total and relative abundances *X* and *π*.

Below, we describe two approaches we propose for sampling *X*, and dealing with peptide uncertainty, based on PEP and FDR filtering, both estimated from proteomics tools such as *MetaMorpheus, Percolator* or *MaxQuant*, and taken as input from *IsoBayes*.

### 2.3 PEP 2-layer latent variable approach

Assume that *N* peptides are detected in total, and that *PEP*_*i*_ is the estimated probability that the *i*-th peptide, albeit absent, is mistakenly detected. First, we sample if a peptide has been erroneously detected, via a Bernoulli distribution:

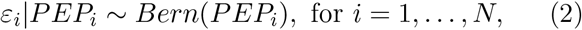

where *ε*_*i*_ = 1 if the *i* − *th* peptide has been mistakenly detected, and 0 if it has been correctly detected. Second, for peptides which are sampled as correctly detected, we spread their abundance to the protein(s) they are compatible with. In particular, we define *Y*_*i*_ as the abundance of the *i*-th peptide, and *ψ*_*i*_ as the list of protein(s) the *i*-th peptide maps to; we further denote by *X*_*pi*_ the (unknown) abundance of peptide *i* that is associated to protein *p*. The *i*-th peptide can be redistributed to the proteins in *ψ*_*i*_ according to the following multinomial distribution:

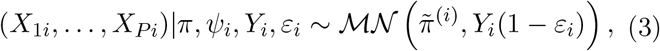

where 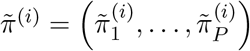, with

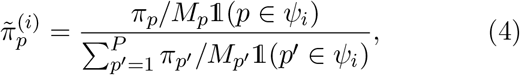

where *M*_*p*_ indicates the number of overall (unique and shared) detected peptides associated to protein isoform *p*, and 𝟙(*A*) is 1 if condition *A* is true, and 0 if condition *A* is false. In other words, 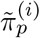 is proportional to *π /M* if the *i*-th peptide maps to the *p*-th protein, and is 0 otherwise. Dividing by *M*_*p*_ ensures that we normalize for the number of peptides contributing to each protein’s abundance. Note that *M*_*p*_ ≥ 1, for *p* = 1, … , *P* , because only isoforms with at least 1 detected peptide are analyzed.

The denominator in (4) ensures that 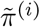 is a probability vector adding to 1; i.e., 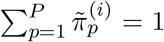. In (3), the peptide abundance, *Y*_*i*_(1 − *ε*_*i*_), is 0 if the peptide has been sampled as mistakenly detected in (2) (i.e., when *ε*_*i*_ = 1). The protein isoform abundances are then recovered by adding the abundances obtained from the *N* peptide allocations: *X*_*p*_ = *X*_*p*1_ + … *X*_*pN*_ , for *p* = 1, … , *P* .

### 2.5 FDR 1-layer latent variable approach

Alternatively, *IsoBayes* can filter peptides with an FDR below a user-defined threshold (usually 0.01). The abundance of each selected peptide is then allocated to the protein(s) it is compatible with, as in (3), with *ε*_*i*_ set to 0 for all peptides *i* = 1, … , *N* . This approach results in a faster runtime, at the cost of a small loss of performance, due to a less accurate propagation of the uncertainty of peptide detections (see results below).

### 2.5 Informative prior

We use a conjugate Dirichlet prior for *π*:

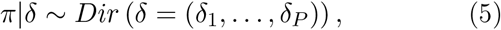

which results in a convenient Dirichlet posterior distribution:

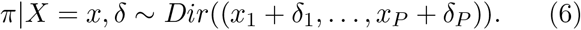

The hyper-parameters *δ* are set proportional to the corresponding relative transcript isoform abundances (informative prior), when available, or to 1 (weakly informative prior), if mRNA data is absent (for more details see Supplementary Details).

### 2.6 Inference

Parameters and latent states are alternately sampled from their conditional distributions, via a Markov chain Monte Carlo (MCMC) scheme, according to two Gibbs samplers (Geman and Geman, 1984; Gelfand and Smith, 1990b): *π*|*X* as in (6), and *X*|*Y, π* as in (2)-(3). Albeit our scheme involves several parameters and latent states, they are all updated via Gibbs samplers, which results in good mixing and convergence. By default, the MCMC is run for 2,000 iterations, with a burn-in of 1,000 iterations (parameters can be increased by users).

The posterior chains of *X* and *π* are then used to compute the output of *IsoBayes*. First, the probability that the *p*-th protein isoform is present, which is given by *Pr* (*X*_*p*_ *>* 0), is estimated as the average time the posterior chain of *X*_*p*_ is positive. Second, estimates of the overall and relative abundances are obtained as the posterior means of *X*_*p*_ and *π*_*p*_, and are accompanied by the respective 0.95 level highest posterior density credible intervals (CIs). Finally, when available, RNA-seq data are used to estimate the relative abundances of transcripts, denoted by 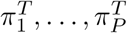. These quantities allow us to compute the log2-FC between protein isoform and transcript relative abundances, i.e., *log*2 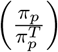, and the probability that the relative abundance is higher in proteins than in transcripts, i.e., *Pr* 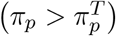, which is computed as the average times the posterior chain of *π*_*p*_ is greater than 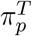 .

Although full MCMC approaches can be computationally intensive, our algorithm is coded in C++, which is a compiled programming language that allows for massive computational gains compared to base R; furthermore, where multiple cores are available, we separate protein isoforms in blocks that have no peptides in common with other blocks, and analyze them in parallel. This allows our algorithm to run within a few minutes (see results below).

## 3 Results

### 3.1 Real data

We collected data dependent acquisition, label-free, liquid chromatography MS data from the *jurkat* and *WTC-11* cell lines, which for simplicity, from now on, we will call *jurkat* and *WTC-11*. The *jurkat* data was collected via Thermo Scientific LTQ Orbitrap Velos mass spectrometer, while the *WTC-11* data was analyzed via Thermo Scientific Orbitrap Eclipse Tribrid mass spectrometer. The *jurkat* and *WTC-11* datasets were collected via 6 (ArgC, AspN, Chym, GluC, LysC, and Trypsin) and 4 (AspN, Chym, LysC, and Trypsin) distinct proteases, respectively, hence leading to 10 datasets overall; each protease consisted of multiple fractions (Supplementary Table 1). The former and latter cell lines were also associated to transcriptomics data, in the form of paired-end (2 × 200 bp) short-read RNA-seq (Illumina HiSeq 2000), and long-read RNA-seq (PacBio Iso-Seq), respectively; we used *kallisto* (Bray et al., 2016) to align short-reads to a reference transcriptome and quantify transcript abundances. Both MS datasets, and respective RNA-seq data, were made publicly available via figshare at https://figshare.com/projects/IsoBayes_paper_data/183988 (see Availability). Complete experimental details for the *jurkat* and *WTC-11* MS sample preparation, digestion and MS analysis can be found in Miller et al. (2019), and in Supplementary Details, respectively; the corresponding RNA-seq datasets are fully described in Sheynkman et al. (2013), and de Souza et al. (2023), respectively.

**Table 1.** Average results, across the 10 proteases, from the simulation study; full results are available in Supplementary Tables 2 and 3. “Abundance present iso” and “Abundance absent iso” indicate the estimated average abundance for protein isoforms which were actually simulated to be present and absent, respectively.

| Method | AUC | log10-corr | 0.95 CI coverage | Abundance absent iso | Abundance present iso |
| --- | --- | --- | --- | --- | --- |
| *IsoBayes* | 0.92 | 0.87 | 0.97 | 0.67 | 8.31 |
| *IsoBayes_mRNA* | 0.97 | 0.97 | 0.99 | 0.16 | 8.60 |

### 3.2 Simulation study

In order to assess the perfomance of our method, we initially benchmarked it in a simulation study, where we: i) generated artificial data, ii) fit *IsoBayes* to it, and iii) compared our estimates with the original ground truth used to simulate the data (i.e., presence/absence and abundance of protein isoforms). In order to generate realistic simulations, we used abundance estimates, and peptide-protein connections (i.e., *ψ*) obtained from real data. We fit *IsoBayes* to each dataset (in the FDR more, and filtering peptides with FDR > 0.01), using PSM counts and mRNA abundances, and inferred protein isoform abundances; we then used these estimates to generate 10 simulated datasets (1 per protease). In order to simulate peptide-level abundances, we rounded each protein isoform abundance to the closest integer, and randomly allocated it to the peptide(s) compatible with it via a multinomial distribution, with equal probability for each peptide.

We then ran *IsoBayes* on the simulated data, with and without mRNA abundances. Note that, transcript information was not explicitly used to generate the simulated data: RNA-seq data is only indirectly associated with the protein isoform abundances we simulated, because it was employed to obtain the estimated abundances on real data. In particular, the average log10-correlation, across proteases, between mRNA and protein isoform abundances is equal to 0.65. We then calculated several metrics of performance of our results: i) the area under the curve (AUC) for the protein isoform presence/absence, via the estimated probability of presence; ii) the Pearson correlation, on the logarithm with base 10 (log10) scale, between the real and estimated protein isoform overall abundances, and iii) the coverage of the 0.95 level CI for the overall abundance (i.e., the fraction of protein isoforms whose real abundance is contained in the estimated CI); iv) the average abundance, separately for protein isoforms that are present and absent in the ground truth. Note that, in ii), to avoid computing the logarithm of 0, a unit is added to all abundances before computing the logarithm; i.e., log10(abundance + 1). Results for all 10 datasets are reported in Supplementary Tables 2 and 3, while average results are shown in Table 1. In all simulations, *IsoBayes* has high AUCs and log10 correlations, and a CI coverage of at least 0.95; as expected, results further improve when incorporating mRNA abundances (i.e., *IsoBayes_mRNA*). This shows that even noisy mRNA prior information (0.65 log10 correlation at the isoform level) leads to an enhancement of the inferential results. Furthermore, the estimated abundances are significantly higher for isoforms which are actually present in the simulation, than for those that are absent (i.e., 12 and 54 times larger for *IsoBayes* and *IsoBayes_mRNA*, respectively).

### 3.3 Real data applications

While simulations enable assessing the performance of a method based on a ground truth, they fail to capture the full complexity of real data; this is particularly true with proteomics data, which is characterized by a large degree of technical and biological noise. However, method validation is challenging on real data, because a ground truth is missing. Here, we benchmarked our approach on the *jurkat* and *WTC-11* real datasets; to overcome the absence of a ground truth, we used an approach which is conceptually similar to the leave-one-out cross validation: for each dataset, we analyzed one protease at a time, and used all the remaining ones (from the same cell line) to validate results. This is possible because all proteases refer to the same cell line, and are expected to lead to highly coherent results. For the validation, we estimated the presence/absence and abundance of protein isoforms via a subset of reliable peptide identifications: peptides with an FDR below 0.01, and mapping to a single protein isoform (also called unique peptides); these peptides have a small probability of being erroneously detected, and no mapping ambiguity. As a consequence of this choice, in the validation step, we removed protein isoforms which are not associated to any (detected or undetected) unique peptide in the theoretical search database, because it is not possible to validate them with our approach.

MS data were pre-processed via two popular proteomics tools, *MetaMorpheus* (Solntsev et al., 2018) and *Percolator* (Käll et al., 2007; The et al., 2016) (from the *OpenMS* (Röst et al., 2016) toolkit), to obtain peptide-level information (i.e., PSM counts, intensities, FDR, PEP, and the peptide-protein compatibility map). We benchmarked our approach against other popular tools for isoform-level inference; namely, *Fido* (Serang et al., 2010), *PIA* (Uszkoreit et al., 2015), and *EPIFANY* (Pfeuffer et al., 2020). A notable method, the *ProteinProphet* (Nesvizhskii et al., 2003), was not considered here, because of the elevated runtime and low performance it exhibited in recent benchmarks (Pfeuffer et al., 2020). All our competitors filter peptides and PSM counts based on their FDR, and two of them (*Fido* and *EPIFANY* ) are integrated within the *OpenMS* toolkit, and cannot be used with the output of other proteomics tools, such as *MetaMorpheus*. Therefore, for a fair comparison, all methods were benchmarked on the output of *OpenMS* ‘ *Percolator*, and only peptides with an FDR below 0.01 were retained. We computed three metrics to quantify the complexity induced by shared peptides (i.e., peptides that map to multiple protein isoforms); among filtered peptides: i) 79% of the PSM counts are associated to shared peptides, ii) 81% of the peptides are shared, and iii) shared peptides are (on average) compatible with 4.9 distinct isoforms (Supplementary Table 4).

Figure 2 shows the receiver operating characteristic (ROC) curve of each method, while Supplementary Table 5 reports the corresponding area under the curve (AUC). In all datasets, *IsoBayes* displays higher sensitivity and specificity than other methods, even when using MS data only; as expected, when incorporating transcriptomics data, performance further improves.

**Figure 2:**
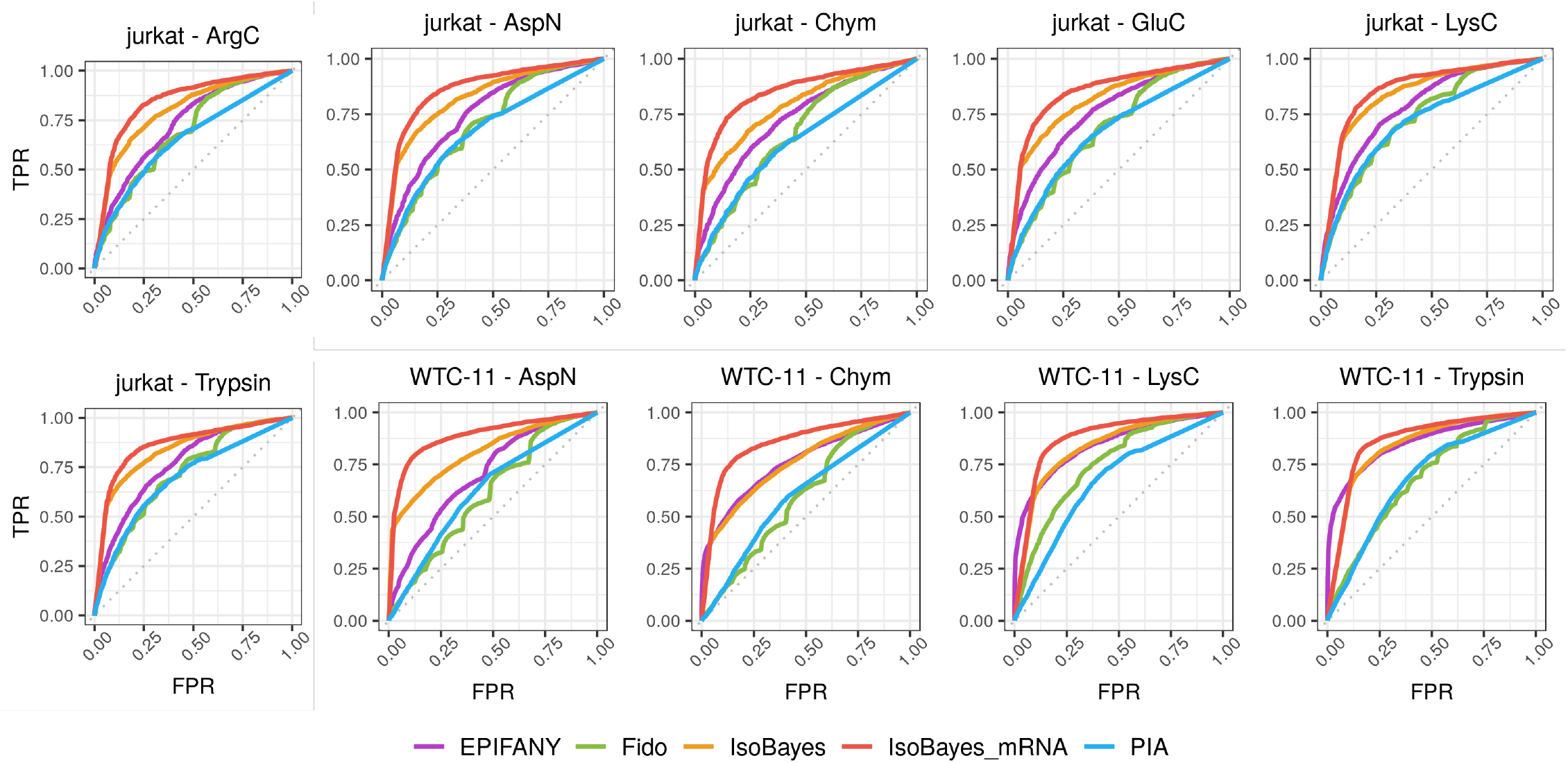
ROC curves, comparing true positive rate (TPR) and false positive rate (FPR), for the identification of protein isoforms in each real dataset.

In addition, to study the quality of our isoform-level abundances, we compared, on the log10 scale (i.e., log10(abundance + 1), our estimates with those detected in our multi-protease validation (Figure 3 and Supplementary Figure 1). To simplify the visual representation, we aggregated isoforms across the proteases from the same cell line. Note that none of our competitors provides abundance measures, therefore here we only evaluated the estimates from our approach. When using MS data only, the log10 correlation is of 0.64 and 0.53, for the *jurkat* and *WTC-11* datasets, respectively; these values increase to 0.71 and 0.69, when embedding mRNA data (Supplementary Table 6). Notably, the mRNA prior leads to a bigger improvement of both AUC and log10 correlation in the *WTC-11* dataset (which uses long-read RNA-seq), compared to the *jurkat* dataset (that uses short-read RNA-seq) (Supplementary Tables 5-6). Although results are not fully comparable, because short and long read protocols are used on distinct datasets, it appears that long-read transcriptomics data can enhance our inference more than short-read data. In the Figures one can see that the scale of our estimates and of the validated abundances do not align; this is because the two are not fully comparable: while our estimates refer to all peptides from a single protease, the validated abundances refer only to unique peptides, but from all proteases (except the one used to compute the estimates). In our simulations, we see that *IsoBayes* provides unbiased abundance estimates: the average difference between estimated and real abundances is −2 × 10^−18^ and 2 × 10^−17^, with and without mRNA abundance, respectively.

**Figure 3:**
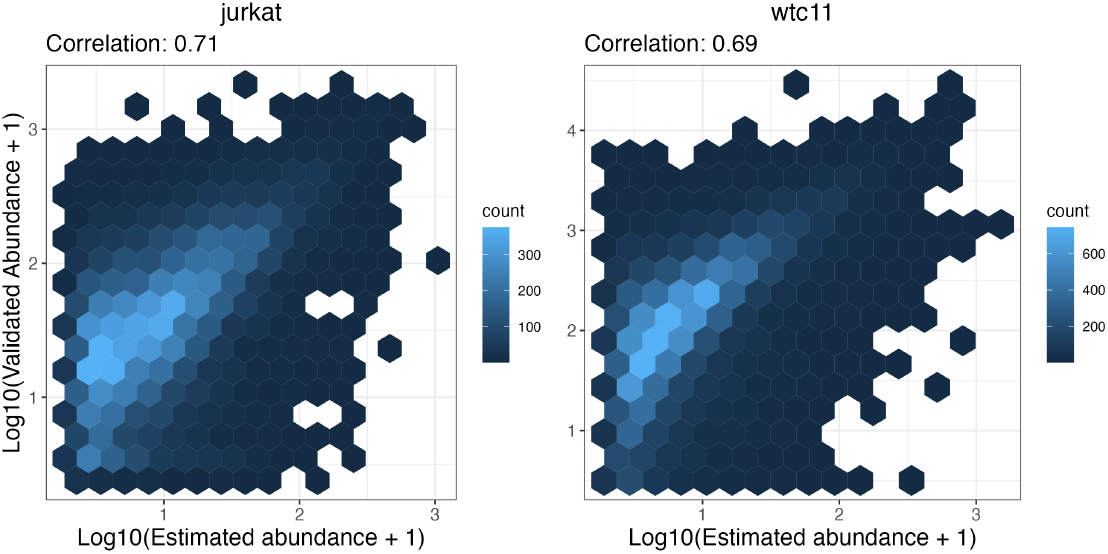
Hexbin plot for the log10 protein isoform abundances (i.e., log10(abundance + 1)), estimated from *IsoBayes_mRNA* (x axis), and found in the validation set (y axis). In each cell line, we considered results from all proteases. Left: *jurkat* dataset; right: *WTC-11* dataset.

We also investigated aggregated results at the gene-level: correlations are higher than at the isoform-level (between 0.87 and 0.89), which is expected given the lower level of (biological and technical) noise at the gene level (Supplementary Table 7, and Supplementary Figures 2-3). Interestingly, while mRNA abundance significantly improves results at the isoform-level, it only marginally impacts inference at the gene-level. This is because, while mRNA data aims at improving the allocation of shared peptides, most peptides are shared across isoforms from the same gene, and there is little mapping ambiguity between distinct genes. In particular, after FDR filtering (0.01 threshold), while on average (across proteases) 71% of PSM counts are shared across multiple isoforms, only 31% of PSM counts are associated to distinct genes (Supplementary Table 4).

We additionally studied our comparison between isoform protein and transcript abundances. Note that, to avoid infinite values, log2-FCs are stabilized by adding a small constant, *κ* = 1.5 × 10^−^6 to both relative abundances; i.e., for each protein isoform *p*, we compute the log2-FC as 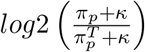. For each protein isoform, we compared the estimated log2-FC with the one obtained in the validation set (i.e., the remaining proteases): results are highly coherent, with correlation values between 0.80 and 0.95 (Supplementary Table 6). In general, mRNA and protein isoform abundances are positively correlated: the correlation of log10 abundances is between 0.45 and 0.66 in our benchmarks (Supplementary Table 8); this supports the usage of mRNA abundance as informative prior in our approach. Nonetheless, there are isoforms, which we aim to detect, where mRNA and protein abundances differ; this is possible because the posterior distribution of *π*, although informed by the mRNA data via the prior, is dominated by the peptide-level data. Indeed, when considering the extreme estimated probabilities 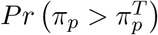 (i.e., below 0.01 and above 0.99), we observe a clear separation of the log2-FCs obtained in the validation set, that we refer to as validated log2-FCs (Figure 4, and Supplementary Figures 4-6). In particular, probabilities near 0 are associated to small values of the validated log2-FC, while probabilities close to 1 lead to large validated log2-FCs; interestingly, probabilities near 0.5 are associated to validated log2-FCs around 0 (i.e., similar relative abundance between proteins and transcripts). Our findings suggest that *IsoBayes* could be helpful in identifying isoforms where protein and transcript relative abundances significantly differ, possibly indicating post translational changes, or different translation rates between isoforms.

**Figure 4:**
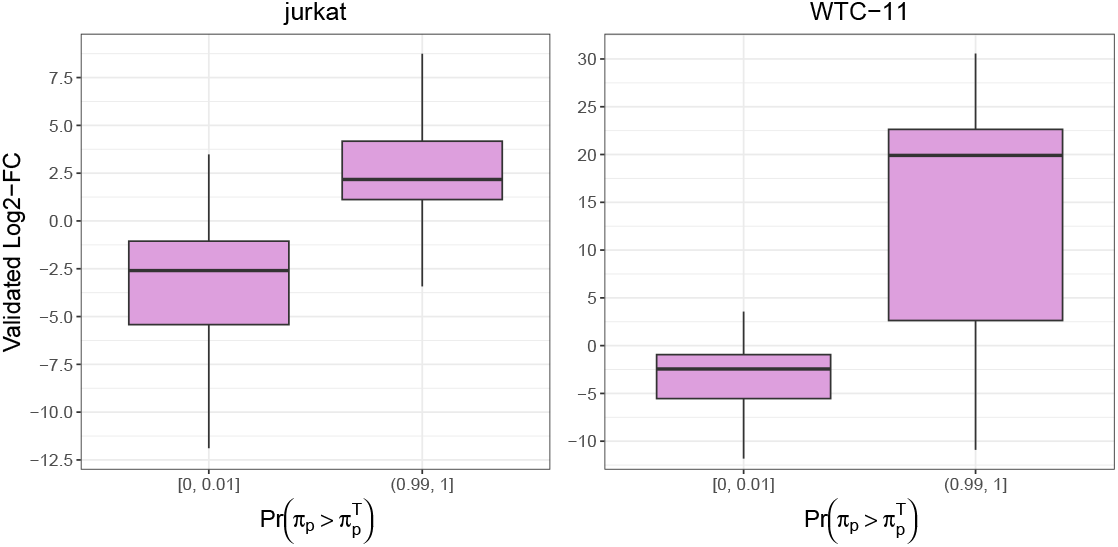
Boxplot of the stabilized log2-FCs between protein and transcript relative abundances, identified in the validated set, stratified based on the probability, estimated by *IsoBayes_mRNA*, that isoform relative abundances are higher at the protein-level than at the transcript-level. Small estimated probabilities (below 0.01) are mainly associated to negative log2-FCs in the validation set; conversely, large estimated probabilities (above 0.99) typically lead to positive log2-FCs in the validation set. In each cell line, we considered results from all proteases. Left: *jurkat* dataset; right: *WTC-11* dataset.

From a computational perspective, the runtimes of all methods are comparable, except for *Fido*, which required significantly more time (Figure 5); conversely, in terms of memory, *EPIFANY* and *Fido* are the most efficient tools, followed by *IsoBayes*, while *PIA* stands as the more expensive approach (Supplementary Figure 7). Finally, we considered the subset of multi-isoform genes (i.e., genes with 2 or more protein isoforms associated to at least one detected peptide), and found that, on average across our datasets, 73% of the abundance estimated from *IsoBayes* is associated to the dominant protein isoform (i.e., the most abundant protein isoform within a gene), while the remaining 27% is associated to non-dominant isoforms. This indicates how much information can be gained when performing inference on isoform-level data, compared to aggregating data at the gene-level.

**Figure 5:**
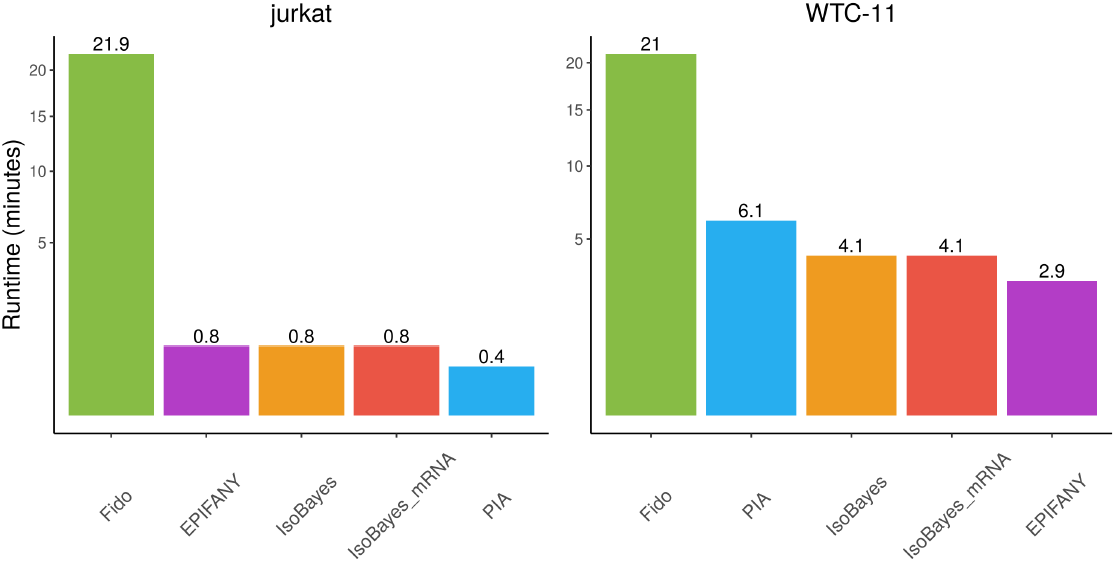
Average runtime of each method, expressed in minutes, across the proteases of a cell-line. Each method was run on 8 cores. Left: *jurkat* dataset; right: *WTC-11* dataset.

### 3.4 Isoforms without unique peptides

In order to investigate the extent to which unique peptides influence results, we also studied the subset of protein isoforms solely associated to shared peptides; these isoforms are the hardest ones to infer, because they are not supported by any detected unique peptide. Although all methods display lower performance, the relative ranking of methods is unchanged, and our estimated abundances (at both isoform- and gene-level) correlate well with our multi-protease validation values (Supplementary Figures 8-10, and Supplementary Table 9). Importantly, the gap between *IsoBayes_mRNA* and the other approaches (including *IsoBayes*) significantly increases; this indicates that incorporating mRNA abundance is particularly beneficial for studying protein isoforms not associated to detected unique peptides. Indeed, it is expected that, in absence of unique peptides, the mRNA-based informative prior plays a crucial role in enhancing the allocation of the abundance from shared peptides. Furthermore, our estimated log2-FCs, between protein and mRNA relative abundances, highly correlate with those estimated in the validation set (Supplementary Table 6), and again our estimates of 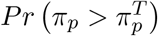 allow a good separation of the log2-FCs in the validation set (Supplementary Figures 11-14), even when mRNA abundance is absent. We also considered isoforms from multi-isoform genes, and found similar results (Supplementary Table 10).

Overall, these results show how *IsoBayes* can be used to infer presence and abundance of protein isoforms, even when unique peptides are absent or undetected; this is particularly beneficial when studying processes such as alternative splicing, where multiple isoforms within a gene are expressed, and peptides are typically shared across the isoforms of a gene.

### 3.5 Robustness to input data

We also tested how robust our results are when varying the input data; in particular, we used *MetaMorpheus*’ proteomics tool to obtain peptide-level abundances. We then fit our approach, with and without mRNA abundances, to both PSM counts and intensities, and compared results to those obtained using *Percolator*. All results are highly coherent, both across proteomics tools and between PSM counts and intensities (Supplementary Figures 15-17, and Supplementary Table 11), which shows that *IsoBayes* outputs are robust with respect to the input data provided.

### 3.6 PEP *vs*. FDR mode

Finally, we tested our PEP mode and compared it to the FDR mode. For this purpose, we used both PSM counts and intensities obtained via *MetaMorpheus*. In the FDR model, we used the classical 0.01 threshold (as in the analyses shown above). Instead, in the PEP mode we used a weak FDR filter. In particular, we removed peptides with an FDR > 10%, because these peptides are associates to an average error probability (i.e., PEP) above 0.95 in all proteases (0.98, on average), and therefore were considered too unreliable to be analyzed; conversely, peptides with an FDR below 0.1 are associated to an average PEP below 0.2 in all proteases (0.13, on average). In all datasets, the PEP mode leads to a small but consistent increase of performance, for both the presence/absence of protein isoforms, and their abundance (Supplementary Table 12). These results are expected, given that the PEP mode allows a better propagation of the peptide detection uncertainty; furthermore, since it considers more peptides (on average, 20% more in our data), it also enables studying a larger number of protein isoforms. At the same time though, because more peptides are analyzed and the latent variable model has an additional layer that needs to be sampled, the PEP model also leads to a significant increase of the runtime, which in our case was between 2 and 10 times, while memory usage only marginally increases (Supplementary Table 13).

## 4 Discussion

In this manuscript, we have presented *IsoBayes*, a novel method for performing isoform-level inference from MS proteomics data. Our approach accounts for the uncertainty in peptide-to-protein mappings, and in peptide detections (in the PEP mode), via a latent variable approach. When available, we allow integrating mRNA data, from short- or long-read RNA-seq technologies, which enhance the accuracy of the allocation of the abundance from shared peptides, and, therefore improve our inferential results. Our method can infer the presence/absence and abundance of individual isoforms, and allows studying changes between protein and transcript relative abundances.

We designed a simulation study, and two real data analyses, based on a multi-protease approach, where we benchmarked our method to state-of-the-art competitors. Our tool displays good sensitivity and specificity when detecting protein isoforms, its abundance estimates highly correlate with those found in the other proteases, and can detect changes between protein and transcript relative abundances. As expected, the availability of mRNA data improves the accuracy of all our results.

Our model performs well even when only considering the subset of protein isoforms without any detected unique peptide. This indicates that *IsoBayes* can be used to investigate processes such as alternative splicing, where peptides are typically shared across multiple isoforms from the same gene.

We also found that our results are robust and consistent across proteomics tools (*MetaMorpheus*, and *Percolator* ), and abundance estimates (PSM counts, and intensities). Finally, we showed that considering the error probability of each peptide (i.e., PEP) allows for a better propagation of the peptide detection uncertainty, compared to using a crude FDR cutoff, and leads to a marginal increase in performance, at the cost of an increased runtime.

Overall, we believe that our tool may be of great utility to life scientists and computational biologists, who aim to investigate protein usage at the isoform level. *IsoBayes* is freely distributed as an R Bioconductor package, and is accompanied by an example vignette and a plotting function; this simplifies its usage, distribution and integration with other bioinformatics and proteomics pipelines.

We would also like to acknowledge some of the limitations of our work. First, while *IsoBayes* FDR mode is computationally on par with competitors, the PEP mode is more computationally intensive, which may be problematic in large datasets. Second, while we account for two major sources of measurement noise (i.e., shared peptides mapping to multiple isoforms, and erroneously detected peptides), other sources of technical uncertainty are neglected, such as missing peptides, peptide detectability levels (i.e., distinct peptides have a different likelihood of being detected), and inaccurate protein reference database. Third, our validation relies on a subset of reliable peptides (uniquely associated to an isoform, and with an FDR below 0.01); while we believe this is a reasonable and reliable approach, we are aware that a proper ground truth is missing.

Finally, we conclude with a look at future perspectives. In our view, this work is not a stand-alone project, but rather lays the foundations for the development of future methods for proteomics/proteogenomics inference at the isoform level. In particular, we aim to extend *IsoBayes* to embed multiple samples (i.e., biological replicates) within a Bayesian hierarchical framework. This will enable performing differential testing between experimental conditions (e.g., healthy *vs*. diseased, or treated *vs*. untreated) at the isoform-level, hence identifying changes, across groups, in protein isoform abundance, and alternative splicing patterns. A second extension concerns the application to single-cell proteomics data. In this case, we aim to investigate how protein isoform abundances vary across cell types, and study the fraction of cells each isoform is detected in.

## Acknowledgements

This work was supported by NIGMS grant R01GM147653 to MRS and LMS, EU-ESF grant to JB. *CMP* ^3^*V dA* is co-founded by European Social Fund, European Regional Development Fund, the Autonomous Region of the Aosta Valley, and the Italian Ministry of Labour and Social Policy (CUP B68H19005520007). We acknowledge the HPC at the data center of Engineering D.HUB in Pont-Saint-Martin.

## Availability

*IsoBayes* is freely available as a Bioconductor R package at: https://bioconductor.org/packages/IsoBayes. All scripts used for our analyses are available on GitHub (https://github.com/SimoneTiberi/IsoBayes_manuscript version v2) and Zenodo (DOI: 10.5281/zenodo.10203419). The *jurkat* and *WTC-11* MS and RNA-seq datasets are available in figshare at https://figshare.com/projects/IsoBayes_paper_data/183988.

## Author contributions

ST and GMS designed the study. ST conceived the statistical method, and wrote the manuscript. ST and JB implemented the method in R, and ran all the analyses. MRS, BTJ, RM, EJ, and GMS collected the real data. CD contributed ideas to the method’s concept. All Authors contributed to editing the manuscript, and approve it.

## References

N. L. Bray, H. Pimentel, P. Melsted, and L. Pachter. Near-optimal probabilistic rna-seq quantification. Nature biotechnology, 34(5):525–527, 2016.

K. Bryan, M.-A. Jarboui, C. Raso, M. Bernal-Llinares, B. McCann, J. Rauch, K. Boldt, and D. J. Lynn. Hiquant: rapid postquantification analysis of large-scale ms-generated proteomics data. Journal of Proteome Research, 15(6):2072–2079, 2016.

B. C. Carlyle, R. R. Kitchen, J. Zhang, R. S. Wilson, T. T. Lam, J. S. Rozowsky, K. R. Williams, N. Sestan, M. B. Gerstein, and A. C. Nairn. Isoform-level interpretation of high-throughput proteomics data enabled by deep integration with rna-seq. Journal of proteome research, 17(10):3431–3444, 2018.

J. Cox and M. Mann. Maxquant enables high peptide identification rates, individualized ppb-range mass accuracies and proteome-wide protein quantification. Nature biotechnology, 26(12):1367–1372, 2008.

R. de Sousa Abreu, L. O. Penalva, E. M. Marcotte, and C. Vogel. Global signatures of protein and mRNA expression levels. Molecular BioSystems, 5(12):1512– 1526, 2009.

V. B. de Souza, B. T. Jordan, E. Tseng, E. A. Nelson, K. K. Hirschi, G. Sheynkman, and M. D. Robinson. Transformation of alignment files improves performance of variant callers for long-read RNA sequencing data. Genome Biology, 24(1):91, 2023.

I. W. Deveson, M. E. Brunck, J. Blackburn, E. Tseng, T. Hon, T. A. Clark, M. B. Clark, J. Crawford, M. E. Dinger, L. K. Nielsen, et al. Universal alternative splicing of noncoding exons. Cell Systems, 6(2):245–255, 2018.

F. Edfors, F. Danielsson, B. M. Hallström, L. Käll, E. Lundberg, F. Pontén, B. Forsström, and M. Uhlén. Gene-specific correlation of rna and protein levels in human cells and tissues. Molecular systems biology, 12(10):883, 2016.

A. E. Gelfand and A. F. Smith. Sampling-based approaches to calculating marginal densities. Journal of the American statistical association, 85(410):398–409, 1990a.

A. E. Gelfand and A. F. Smith. Sampling-based approaches to calculating marginal densities. Journal of the American statistical association, 85(410):398–409, 1990b.

S. Geman and D. Geman. Stochastic relaxation, gibbs distributions, and the bayesian restoration of images. IEEE Transactions on pattern analysis and machine intelligence, (6):721–741, 1984.

L. Käll, J. D. Canterbury, J. Weston, W. S. Noble, and M. J. MacCoss. Semi-supervised learning for peptide identification from shotgun proteomics datasets. Nature methods, 4(11):923–925, 2007.

Y. Liu, A. Beyer, and R. Aebersold. On the dependency of cellular protein levels on mRNA abundance. Cell, 165(3):535–550, 2016.

Y. Liu, M. Gonzàlez-Porta, S. Santos, A. Brazma, J. C. Marioni, R. Aebersold, A. R. Venkitaraman, and V. O. Wickramasinghe. Impact of alternative splicing on the human proteome. Cell reports, 20(5):1229–1241, 2017.

P. Lu, C. Vogel, R. Wang, X. Yao, and E. M. Marcotte. Absolute protein expression profiling estimates the relative contributions of transcriptional and translational regulation. Nature biotechnology, 25(1):117–124, 2007.

C. Ma, S. Xu, G. Liu, X. Liu, X. Xu, B. Wen, and S. Liu. Improvement of peptide identification with considering the abundance of mRNA and peptide. BMC bioinformatics, 18:1–8, 2017.

W.-T. Ma, Z.-Y. Liu, X.-Z. Chen, Z.-L. Lin, Z.-B. Zheng, W.-G. Miao, and S.-Q. Xie. A protein identification algorithm for tandem mass spectrometry by incorporating the abundance of mRNA into a binomial probability scoring model. Journal of proteomics, 197:53–59, 2019.

T. Maier, M. Güell, and L. Serrano. Correlation of mRNA and protein in complex biological samples. FEBS letters, 583(24):3966–3973, 2009.

R. M. Miller, R. J. Millikin, C. V. Hoffmann, S. K. Solntsev, G. M. Sheynkman, M. R. Shortreed, and L. M. Smith. Improved protein inference from multiple protease bottom-up mass spectrometry data. Journal of proteome research, 18(9):3429–3438, 2019.

R. M. Miller, B. T. Jordan, M. M. Mehlferber, E. D. Jeffery, C. Chatzipantsiou, S. Kaur, R. J. Millikin, Y. Dai, S. Tiberi, P. J. Castaldi, et al. Enhanced protein isoform characterization through long-read proteogenomics. Genome biology, 23(1):1–28, 2022.

I. Nesvizhskii, A. Keller, E. Kolker, and R. Aebersold. A statistical model for identifying proteins by tandem mass spectrometry. Analytical chemistry, 75 (17):4646–4658, 2003.

J. Pfeuffer, T. Sachsenberg, T. M. Dijkstra, O. Serang, K. Reinert, and O. Kohlbacher. Epifany: a method for efficient high-confidence protein inference. Journal of proteome research, 19(3):1060–1072, 2020.

S. R. Ramakrishnan, C. Vogel, J. T. Prince, R. Wang, Z. Li, L. O. Penalva, M. Myers, E. M. Marcotte, and D. P. Miranker. Integrating shotgun proteomics and mRNA expression data to improve protein identification. Bioinformatics, 25 (11):1397–1403, 2009.

H. L. Röst, T. Sachsenberg, S. Aiche, C. Bielow, H. Weisser, F. Aicheler, S. Andreotti, H.-C. Ehrlich, P. Gutenbrunner, E. Kenar, et al. Openms: a flexible open-source software platform for mass spectrometry data analysis. Nature methods, 13(9):741–748, 2016.

Salovska, H. Zhu, T. Gandhi, M. Frank, W. Li, G. Rosenberger, C. Wu, P.L. Germain, H. Zhou, Z. Hodny, et al. Isoform-resolved correlation analysis between mRNA abundance regulation and protein level degradation. Molecular systems biology, 16(3):e9170, 2020.

O. Serang, M. J. MacCoss, and W. S. Noble. Efficient marginalization to compute protein posterior probabilities from shotgun mass spectrometry data. Journal of proteome research, 9(10):5346–5357, 2010.

G. M. Sheynkman, M. R. Shortreed, B. L. Frey, and L. M. Smith. Discovery and mass spectrometric analysis of novel splice-junction peptides using rna-seq. Molecular & Cellular Proteomics, 12(8):2341–2353, 2013.

S. K. Solntsev, M. R. Shortreed, B. L. Frey, and L. M. Smith. Enhanced global post-translational modification discovery with MetaMorpheus. Journal of proteome research, 17(5):1844–1851, 2018.

M. A. Tanner and W. H. Wong. The calculation of posterior distributions by data augmentation. Journal of the American statistical Association, 82(398):528– 540, 1987.

M. The, M. J. MacCoss, W. S. Noble, and L. Käll. Fast and accurate protein false discovery rates on large-scale proteomics data sets with percolator 3.0. Journal of the American Society for Mass Spectrometry, 27:1719–1727, 2016.

J. Uszkoreit, A. Maerkens, Y. Perez-Riverol, H. E. Meyer, K. Marcus, C. Stephan, O. Kohlbacher, and M. Eisenacher. Pia: an intuitive protein inference engine with a web-based user interface. Journal of proteome research, 14(7):2988– 2997, 2015.

A. Vesvizhskii and A. Aebersold. Interpretation of shotgun proteomic data: the protein interference problem. Mol Cell Proteomics, 4:1419–40, 2005.

D. Wang, B. Eraslan, T. Wieland, B. Hallström, T. Hopf, D. P. Zolg, J. Zecha, A. Asplund, L.-h. Li, C. Meng, et al. A deep proteome and transcriptome abundance atlas of 29 healthy human tissues. Molecular systems biology, 15 (2):e8503, 2019.

C. Willyard. Expanded human gene tally reignites debate. Nature, 558(7710):354–355, 2018.

