## Supplementary material for "*IsoBayes*: a Bayesian approach for single-isoform proteomics inference" (from another version of this preprint: version 2, 2025-04-30)

[1]*Computational and Chemical Biology, Italian Institute of Technology, CMP[3]VdA, Aosta, Italy.*
[2]*Astronomical Observatory of the Autonomous Region of the Aosta Valley (OAVdA), Nus, Italy.*
[3]*Department of Chemistry, University of Wisconsin-Madison, Madison, WI, USA.*
[4]*Frederick National Laboratory for Cancer Research, Frederick, MD, USA.*
[5]*Department of Molecular Physiology and Biological Physics, University of Virginia, Charlottesville, VA, USA.*
[6]*Centre Européen de Calcul Atomique et Moléculaire, École Polytechnique Fédérale de Lausanne, Lausanne, Switzerland.*
[7]*Department of Biostatistics and Medical Informatics, University of Wisconsin-Madison, Madison, WI, USA.*
[8]*Department of Statistical Sciences, University of Bologna, Bologna, Italy.*
[*]

#### 1 Supplementary Details

##### 1.1 Mathematical details

#### Y | X

Our method, as illustrated in the graphical model in Figure 1 of the main text, involves hyper-parameters $\delta$, parameters $\pi$, latent variables $X$ (i.e., protein-isoform abundances), and observations $Y$ (i.e., peptide-level abundances). In the main paper, we define the conditional distributions of $\pi|\delta$, and $X|\pi$, in formulae (5) and (1), respectively. Below, we describe how $Y$ is derived from $X$. Let $i_1, \dots, i_{M_p}$ be the indices of the $M_p$ detected peptides associated to protein isoform $p$; then:

$$\left(Z_{p,i_1}, \dots, Z_{p,i_{M_P}}\right)\Big| X_p, \sim \mathcal{MN}\left(\frac{1}{M_p}, n = X_p\right), \text{ for } p = 1, \dots, P, \tag{1}$$

where $Z_{p,i}$ indicates the abundance of protein $p$ associated to its peptide $i$. The overall abundance of peptide $i$ can then be deterministically obtained as $Y_i = \sum_{p \in \psi_i} Z_{p,i}$, for $i = 1, \dots, N$.

###### Informative prior

*A priori*, we assume that $\pi$, the relative abundance of protein isoforms, follows a Dirichlet distribution with parameter $\delta = (\delta_1, \dots, \delta_P)$:

$$\pi|\delta \sim Dir(\delta), \tag{2}$$

which results in the following posterior distribution:

$$\pi|X = x, \delta \sim Dir((x_1 + \delta_1, \dots, x_P + \delta_P)). \tag{3}$$

If mRNA is absent, we use a weakly informative prior and set $\delta_1 = \dots = \delta_P = 1$.
Instead, when mRNA data is available, $\delta$ is (almost) proportional to the mRNA isoform relative abundance, estimated from transcriptomics data $pi^T = \left(\pi_1^T, \dots, \pi_P^T\right)$. In particular, in order to avoid setting prior values to 0,

we add a small constant, $\epsilon = 10^{-5}$, to the values in $pi^T$, and obtain $\tilde{\pi}_p^T = \pi_p^T + \epsilon$, for $p = 1, \ldots, P$. We re-normalize these values to make sure they add to 1: $\hat{\pi}_p^T = \frac{\tilde{\pi}_p^T}{\sum_{p'=1}^{P} \tilde{\pi'}_p^T}$. Then, we compute the overall abundance in the data (PSM counts or intensities): $x_{TOT} = x_1 + \ldots,_P$. Finally, we set $\delta_p = \hat{\pi}_p^T \times c \times x_{TOT}$, for $p = 1, \ldots, P$, with $c = 0.1$. In other words, $\delta_p$ is proportional to $\hat{\pi}_p^T$, with weight denoted by $c \times x_{TOT}$, where $c$ is a constant that denotes how weight the prior has in the posterior calculation. Note that $c$ is not a tuning parameter, and is kept fixed at 0.1. This value implies that, in the posterior distribution in (3), the prior globally weights 10 times less than the data (i.e., protein abundances): $\sum_{p=1}^{p} \delta_p = 0.1 x_{TOT}$, while $\sum_{p=1}^{p} x_p = 0.1 x_{TOT}$.
We performed a sensitivity analysis to study how different values of $c$ affect our inference (Supplementary Figure 18), and found that results are highly robust when $c$ is between 0.1 and 0.4, and deteriorates for larger values of $c$ (i.e., then the prior has a strong weight in the posterior). Therefore, we conservatively choose a value of 0.1, which in general provides the highest AUC in our benchmarks.

#### 1.2 WTC-11 mass spectrometry data

##### Mass spectrometry-based proteomics sample preparation

Harvested WTC11 cells (induced pluripotent stem cells from adult skin), approximately 8 million cells in each sample, were pelleted and frozen at -80 celsius degrees. The sample pellets were probe-sonicated and lysed according to the Filter Aided Sample Preparation (FASP) protocol [2]. The Lysis buffer used in the FASP was changed to 6% SDS, 150 mM DTT, 75 mM Tris-HCl. BCA assay of lysates indicated approximately 1.4 $\mu$g/$\mu$L protein concentration. Separate aliquots were digested with either trypsin, chymotrypsin, LysC, or AspN (1 $\mu$g enzyme per digestion) overnight at 37 celsius degrees (chymotrypsin digest was room temperature). The total peptide quantity subjected to subsequence offline HPLC fractionation, as estimated by Nanodrop, was: 85 $\mu$g (trypsin), 130 $\mu$g (LysC), 60 $\mu$g (AspN), and 100 $\mu$g (chymotrypsin).

##### Offline HPLC Fractionation

From each digest, peptides were pooled and dried down to a volume of 40 $\mu$L and subjected to offline high pH RP-HPLC fractionation using an Agilent 1200 HPLC and 96 well plate for fraction collection. Samples were loaded onto a Thermo Scientific Hypersil Gold C18 column (150 mm x 3 mm x 3 µm C18), equilibrated with 95% solvent A (20 mM ammonium formate, pH of 10) and 5% solvent B (70% acetonitrile, and 30% solvent A), and eluted at a flow rate of 400 µL/min, with fractions collected into wells every 30 seconds from RT 30-63 min. The following gradient was used: 5% B from 0-30 min, 5-65% B from 30-63 min, 65-100% B from 64-69 min, 100-5% B from 69-70 min, 5% B from 70-73 min. Approximately 20-30 peptide-containing fractions from each offline fractionation were selected for LC-MS/MS analysis (Supplementary Table 1).

##### NanoLC-MS/MS analysis

The resulting peptides were dried to 12 $\mu$L and analyzed by nanoLC-MS/MS using a Dionex Ultimate 3000 (Thermo Fisher Scientific, Bremen, Germany) coupled to an Orbitrap Eclipse Tribrid mass spectrometer (Thermo Fisher Scientific, Bremen, Germany). Three microliters of each peptide-containing sample were loaded onto an Acclaim PepMap 100 trap column (300 $\mu$m x 5 mm x 5 $\mu$m C18) and gradient-eluted from an Acclaim PepMap 100 analytical column (75 $\mu$m x 25 cm, 3 $\mu$m C18) equilibrated in 96% solvent A (0.1% formic acid in water) and 4% solvent B (80% acetonitrile in 0.1% formic acid). The peptides were eluted at 300 nL/min using the following 4hr gradient (trypsin and LysC samples): 4% B from 0-5 min, 4 to 28% B from 5-210 min, 28-40% B from 210-240 min, 40-95% B from 240-250 min and 95% B from 250-260 min or 2hr gradient (chymo and AspN samples): 4% B from 0-5 min, 4-28% B from 5-100 min, 28-40% B from 100-115 min, 40-95% B from 115-121 min, 95% B from 121-123 min.
The Orbitrap Eclipse was operated in positive ion mode with 1.9 kV at the spray source, RF lens at 30% and data dependent MS/MS acquisition with XCalibur version 4.3.73.11. Positive ion Full MS scans were acquired in the Orbitrap from 375-1500 m/z with 120,000 resolution. Data dependent selection of precursor ions was performed in Cycle Time mode, with 3 seconds in between Master Scans, using an intensity threshold of 2 x 104 ion counts and applying dynamic exclusion (n=1 scans within 30 seconds for an exclusion duration of 60 seconds and +/- 10 ppm mass tolerance). Monoisotopic peak determination was applied and charge states 2-7 were included for HCD scans

(quadrupole isolation mode; 1.6 m/z isolation window). The resulting fragments were detected in the Orbitrap at 15,000 resolution with standard AGC target.

##### MS database search

Standard proteomic analysis of acquired mass spectra files were performed using the free and open source search software program MetaMorpheus [1].

### 2 Supplementary Tables

#### 2.1 Real data

| Cell line | Protease | Number of Fractions |
|---|---|---|
| jurkat | ArgC | 11 |
| jurkat | AspN | 11 |
| jurkat | Chym | 6 |
| jurkat | GluC | 11 |
| jurkat | LysC | 11 |
| jurkat | Trypsin | 10 |
| WTC-11 | AspN | 35 |
| WTC-11 | Chym | 18 |
| WTC-11 | LysC | 30 |
| WTC-11 | Trypsin | 20 |

**Supplementary Table 1:** Number of fractions in each protease, in the real data.

#### 2.2 Simulation study

| Cell line | Protease | AUC | log10-corr | 0.95 CI coverage | Abundance absent iso | Abundance present iso |
|---|---|---|---|---|---|---|
| jurkat | ArgC | 0.91 | 0.86 | 0.98 | 0.69 | 5.73 |
| jurkat | AspN | 0.93 | 0.89 | 0.98 | 0.60 | 8.73 |
| jurkat | Chym | 0.87 | 0.83 | 0.98 | 0.71 | 4.41 |
| jurkat | GluC | 0.93 | 0.90 | 0.98 | 0.59 | 9.14 |
| jurkat | LysC | 0.95 | 0.92 | 0.99 | 0.48 | 11.84 |
| jurkat | Trypsin | 0.95 | 0.92 | 0.98 | 0.47 | 11.30 |
| WTC-11 | AspN | 0.92 | 0.82 | 0.96 | 0.82 | 7.28 |
| WTC-11 | Chymo | 0.89 | 0.78 | 0.94 | 1.16 | 9.62 |
| WTC-11 | LysC | 0.93 | 0.86 | 0.97 | 0.65 | 7.43 |
| WTC-11 | Trypsin | 0.93 | 0.88 | 0.98 | 0.70 | 7.26 |
| Average | | 0.92 | 0.87 | 0.98 | 0.69 | 8.28 |

**Supplementary Table 2:** Summary results, from the simulation study, for *IsoBayes* fit without mRNA abundances. “Abundance present iso” and “Abundance absent iso” indicate the estimated average abundance for protein isoforms which were actually simulated to be present and absent, respectively.

| Cell line | Protease | AUC | log10-corr | 0.95 CI coverage | Abundance absent iso | Abundance present iso |
|---|---|---|---|---|---|---|
| jurkat | ArgC | 0.97 | 0.95 | 0.99 | 0.24 | 6.01 |
| jurkat | AspN | 0.97 | 0.97 | 0.99 | 0.16 | 8.97 |
| jurkat | Chym | 0.96 | 0.96 | 0.99 | 0.16 | 4.91 |
| jurkat | GluC | 0.98 | 0.97 | 0.99 | 0.16 | 9.40 |
| jurkat | LysC | 0.98 | 0.98 | 0.99 | 0.15 | 11.98 |
| jurkat | Trypsin | 0.98 | 0.98 | 0.99 | 0.13 | 11.46 |
| WTC-11 | AspN | 0.97 | 0.96 | 0.99 | 0.17 | 7.62 |
| WTC-11 | Chymo | 0.97 | 0.97 | 1.00 | 0.18 | 10.13 |
| WTC-11 | LysC | 0.97 | 0.97 | 0.99 | 0.17 | 7.68 |
| WTC-11 | Trypsin | 0.96 | 0.96 | 0.99 | 0.18 | 7.46 |
| Average | | 0.97 | 0.97 | 0.99 | 0.17 | 8.56 |

**Supplementary Table 3:** Summary results, from the simulation study, for *IsoBayes* fit with mRNA abundances. "Abundance present iso" and "Abundance absent iso" indicate the estimated average abundance for protein isoforms which were actually simulated to be present and absent, respectively.

#### 2.3 Real data - All isoforms

| Cell line | Protease | Multi counts | Multi peptides | N isoforms per multi peptide | Gene multi counts |
|---|---|---|---|---|---|
| jurkat | ArgC | 0.77 | 0.80 | 4.62 | 0.13 |
| jurkat | AspN | 0.76 | 0.82 | 4.73 | 0.09 |
| jurkat | Chym | 0.77 | 0.81 | 4.82 | 0.11 |
| jurkat | GluC | 0.76 | 0.82 | 4.80 | 0.09 |
| jurkat | LysC | 0.78 | 0.83 | 4.85 | 0.10 |
| jurkat | Trypsin | 0.78 | 0.84 | 5.01 | 0.12 |
| WTC-11 | AspN | 0.82 | 0.81 | 5.05 | 0.63 |
| WTC-11 | Chym | 0.83 | 0.79 | 5.08 | 0.63 |
| WTC-11 | LysC | 0.82 | 0.78 | 4.97 | 0.60 |
| WTC-11 | Trypsin | 0.82 | 0.79 | 5.15 | 0.62 |
| Average | | 0.79 | 0.81 | 4.91 | 0.31 |

**Supplementary Table 4:** Fraction of counts from shared peptides (column "Multi counts"); fraction shared peptides (column "Multi peptides"); average number of protein isoforms associated to shared peptides (column "N isoforms per multi peptide"); fraction of counts from peptides shared across multiple genes (column "Gene multi counts"). We considered peptides that pass the 0.01 FDR threshold. shared peptides are peptides associated to at least 2 protein isoforms.

| Cell line | Protease | *IsoBayes_mRNA* | *IsoBayes* | *EPIFANY* | *PIA* | *Fido* |
|---|---|---|---|---|---|---|
| jurkat | ArgC | 0.84 | 0.80 | 0.73 | 0.65 | 0.69 |
| jurkat | AspN | 0.86 | 0.82 | 0.75 | 0.67 | 0.7 |
| jurkat | Chym | 0.85 | 0.78 | 0.73 | 0.63 | 0.67 |
| jurkat | GluC | 0.85 | 0.81 | 0.76 | 0.67 | 0.69 |
| jurkat | LysC | 0.86 | 0.84 | 0.78 | 0.71 | 0.73 |
| jurkat | Trypsin | 0.86 | 0.84 | 0.77 | 0.69 | 0.71 |
| WTC-11 | AspN | 0.89 | 0.81 | 0.70 | 0.62 | 0.61 |
| WTC-11 | Chym | 0.85 | 0.75 | 0.76 | 0.60 | 0.60 |
| WTC-11 | LysC | 0.88 | 0.84 | 0.84 | 0.67 | 0.75 |
| WTC-11 | Trypsin | 0.87 | 0.84 | 0.85 | 0.68 | 0.68 |
| Average | | 0.86 | 0.81 | 0.77 | 0.66 | 0.68 |

**Supplementary Table 5:** Area under the curve (AUC) for the detection of protein isoforms, for every method in each real dataset.

| Correlation | Method | All isoforms | | No unique peptides | |
|---|---|---|---|---|---|
| | | *jurkat* | *WTC-11* | *jurkat* | *WTC-11* |
| log10-Abundance | IsoBayes_mRNA | 0.71 | 0.69 | 0.63 | 0.61 |
| | IsoBayes | 0.64 | 0.53 | 0.51 | 0.38 |
| log2-FCs | IsoBayes_mRNA | 0.85 | 0.95 | 0.85 | 0.93 |
| | IsoBayes | 0.80 | 0.92 | 0.81 | 0.90 |

**Supplementary Table 6:** Pearson correlation values for: i) log10 estimated protein isoform abundances (i.e., log10(abundance + 1); ii) stabilized log2-FCs between protein and transcript relative abundances. In both cases, the correlation is computed between the values estimated on the protease being analyzed, and those found in the validation set. In each cell line, we considered results from all proteases. "All isoforms" indicates results from all protein isoforms, while "No unique peptides" refers to the subset of protein isoforms solely associated to shared peptides.

| Method | *jurkat* | *WTC-11* |
|---|---|---|
| IsoBayes_mRNA | 0.87 | 0.89 |
| IsoBayes | 0.88 | 0.88 |

**Supplementary Table 7:** Correlation between log10 estimated protein gene abundances (i.e., log10(abundance + 1)), and those found in the validation set. In each cell line, we considered results from all protesease.

| | jurkat | WTC-11 |
|---|---|---|
| IsoBayes | 0.52 | 0.45 |
| IsoBayes_mRNA | 0.65 | 0.66 |

**Supplementary Table 8:** Correlation between log10 mRNA and estimated protein isoform abundances (i.e., log10(abundance + 1)). In each cell line, we considered results from all protesease

#### 2.4 Real data - Isoforms without unique peptides

| Cell line | Protease | *IsoBayes_mRNA* | *IsoBayes* | *EPIFANY* | *PIA* | *Fido* |
|---|---|---|---|---|---|---|
| jurkat | ArgC | 0.81 | 0.73 | 0.67 | 0.55 | 0.66 |
| jurkat | AspN | 0.82 | 0.75 | 0.69 | 0.57 | 0.67 |
| jurkat | Chym | 0.80 | 0.70 | 0.66 | 0.53 | 0.63 |
| jurkat | GluC | 0.80 | 0.73 | 0.68 | 0.57 | 0.65 |
| jurkat | LysC | 0.82 | 0.78 | 0.73 | 0.62 | 0.71 |
| jurkat | Trypsin | 0.81 | 0.76 | 0.71 | 0.60 | 0.69 |
| WTC-11 | AspN | 0.83 | 0.69 | 0.64 | 0.53 | 0.60 |
| WTC-11 | Chym | 0.82 | 0.66 | 0.66 | 0.51 | 0.58 |
| WTC-11 | LysC | 0.85 | 0.75 | 0.72 | 0.57 | 0.69 |
| WTC-11 | Trypsin | 0.83 | 0.77 | 0.72 | 0.60 | 0.69 |
| Average | | 0.82 | 0.73 | 0.69 | 0.56 | 0.66 |

**Supplementary Table 9:** Area under the curve (AUC) for the detection of protein isoforms, for every method in each real dataset, computed on the subset of protein isoforms solely associated to shared peptides.

#### 2.5 Real data - Isoforms from multi-isoform genes

| Cell line | Protease | *IsoBayes_mRNA* | *IsoBayes* | *EPIFANY* | *PIA* | *Fido* |
|---|---|---|---|---|---|---|
| jurkat | ArgC | 0.83 | 0.78 | 0.74 | 0.61 | 0.73 |
| jurkat | AspN | 0.84 | 0.8 | 0.75 | 0.64 | 0.73 |
| jurkat | Chym | 0.81 | 0.74 | 0.7 | 0.59 | 0.68 |
| jurkat | GluC | 0.82 | 0.78 | 0.74 | 0.64 | 0.72 |
| jurkat | LysC | 0.85 | 0.83 | 0.78 | 0.7 | 0.77 |
| jurkat | Trypsin | 0.84 | 0.82 | 0.77 | 0.69 | 0.76 |
| WTC-11 | AspN | 0.86 | 0.75 | 0.7 | 0.59 | 0.64 |
| WTC-11 | Chym | 0.83 | 0.7 | 0.68 | 0.55 | 0.61 |
| WTC-11 | LysC | 0.87 | 0.81 | 0.79 | 0.65 | 0.74 |
| WTC-11 | Trypsin | 0.86 | 0.83 | 0.81 | 0.67 | 0.71 |
| Average | | 0.84 | 0.78 | 0.75 | 0.63 | 0.71 |

**Supplementary Table 10:** Area under the curve (AUC) for the detection of protein isoforms, for every method in each real dataset, computed on the subset of protein isoforms from multi-isoform genes (i.e., genes with more than one expressed isoform).

#### 2.6 Real data - Robustness to input data

| Cell line | Metric | *IsoBayes* | | | *IsoBayes_mRNA* | | |
|---|---|---|---|---|---|---|---|
| | | *OpenMS* | *MM PSM* | *MM int* | *OpenMS* | *MM PSM* | *MM int* |
| jurkat | AUC | 0.79 | 0.81 | 0.81 | 0.84 | 0.86 | 0.86 |
| | Correlation | 0.57 | 0.58 | 0.6 | 0.64 | 0.64 | 0.66 |
| WTC-11 | AUC | 0.78 | 0.81 | 0.81 | 0.86 | 0.88 | 0.88 |
| | Correlation | 0.49 | 0.49 | 0.44 | 0.64 | 0.64 | 0.58 |

**Supplementary Table 11:** Area under the curve (AUC) for the detection of protein isoforms, and correlation between log10 estimated protein isoform abundances (i.e., log10(abundance + 1)), and those found in the validation set. Values represent averages across the proteases of the jurkat and WTC-11 datasets. Results refer to *IsoBayes* and *IsoBayes_mRNA*, computed on three input data: i) PSM counts from *OpenMS*' *Percolator* ("OpenMS" column); ii) PSM counts from *MetaMorpheus* ("MM PSM" column); iii) peptide intensities from *MetaMorpheus*("MM int" column). Note that numbers slightly differ with respect to other Tables; this is because, here, we focus on the isoforms that are in common across the three data types, i.e., with at least 1 detected (shared or unique) peptide. This ensures a fair comparison across inputs.

#### 2.7 Real data - PEP *vs.* FDR mode

| Input data | Cell line | Metric | *IsoBayes* | | *IsoBayes_mRNA* | |
|---|---|---|---|---|---|---|
| | | | *PEP mode* | *FDR mode* | *PEP mode* | *FDR mode* |
| MM PSM | jurkat | AUC | 0.82 | 0.81 | 0.86 | 0.85 |
| | | Correlation | 0.60 | 0.60 | 0.65 | 0.65 |
| | WTC-11 | AUC | 0.81 | 0.81 | 0.88 | 0.87 |
| | | Correlation | 0.51 | 0.51 | 0.63 | 0.64 |
| MM int | jurkat | AUC | 0.82 | 0.81 | 0.87 | 0.86 |
| | | Correlation | 0.62 | 0.61 | 0.68 | 0.67 |
| | WTC-11 | AUC | 0.81 | 0.81 | 0.88 | 0.87 |
| | | Correlation | 0.48 | 0.45 | 0.61 | 0.59 |

**Supplementary Table 12:** Area under the curve (AUC) for the detection of protein isoforms, and correlation between log10 estimated protein isoform abundances (i.e., log10(abundance + 1)), and those found in the validation set. Values represent averages across the proteases of the jurkat and WTC-11 datasets. Results refer to *IsoBayes* and *IsoBayes_mRNA*, based the PEP and FDR modes. Methods were fit on both PSM counts ("MM PSM" rows), and peptide intensities ("MM int" rows), computed via *MetaMorpheus*.

| Input data | Cell line | Metric | *IsoBayes* | | *IsoBayes_mRNA* | |
|---|---|---|---|---|---|---|
| | | | *PEP mode* | *FDR mode* | *PEP mode* | *FDR mode* |
| MM PSM | jurkat | Runtime (mins) | 0.9 | 0.4 | 0.9 | 0.4 |
| | | Memory (GB) | 5.4 | 3.3 | 5.0 | 3.3 |
| | WTC-11 | Runtime (mins) | 4.1 | 0.7 | 4.1 | 0.7 |
| | | Memory (GB) | 5.7 | 3.5 | 6.5 | 5.2 |
| MM int | jurkat | Runtime (mins) | 0.9 | 0.4 | 0.9 | 0.4 |
| | | Memory (GB) | 5.5 | 3.3 | 5.9 | 3.4 |
| | WTC-11 | Runtime (mins) | 4.4 | 0.6 | 4.3 | 0.7 |
| | | Memory (GB) | 6.4 | 3.9 | 6.5 | 4.9 |

**Supplementary Table 13:** Runtime (in minutes) and memory (in Gigabytes) for *IsoBayes* and *IsoBayes_mRNA*, based the PEP and FDR modes. Values represent averages across the proteases of the jurkat and WTC-11 datasets. Methods were fit on both PSM counts ("MM PSM" rows), and peptide intensities ("MM int" rows), computed via *MetaMorpheus*.

### 3 Supplementary Figures

#### 3.1 All protein isoforms

##### 3.1.1 Isoform-level results

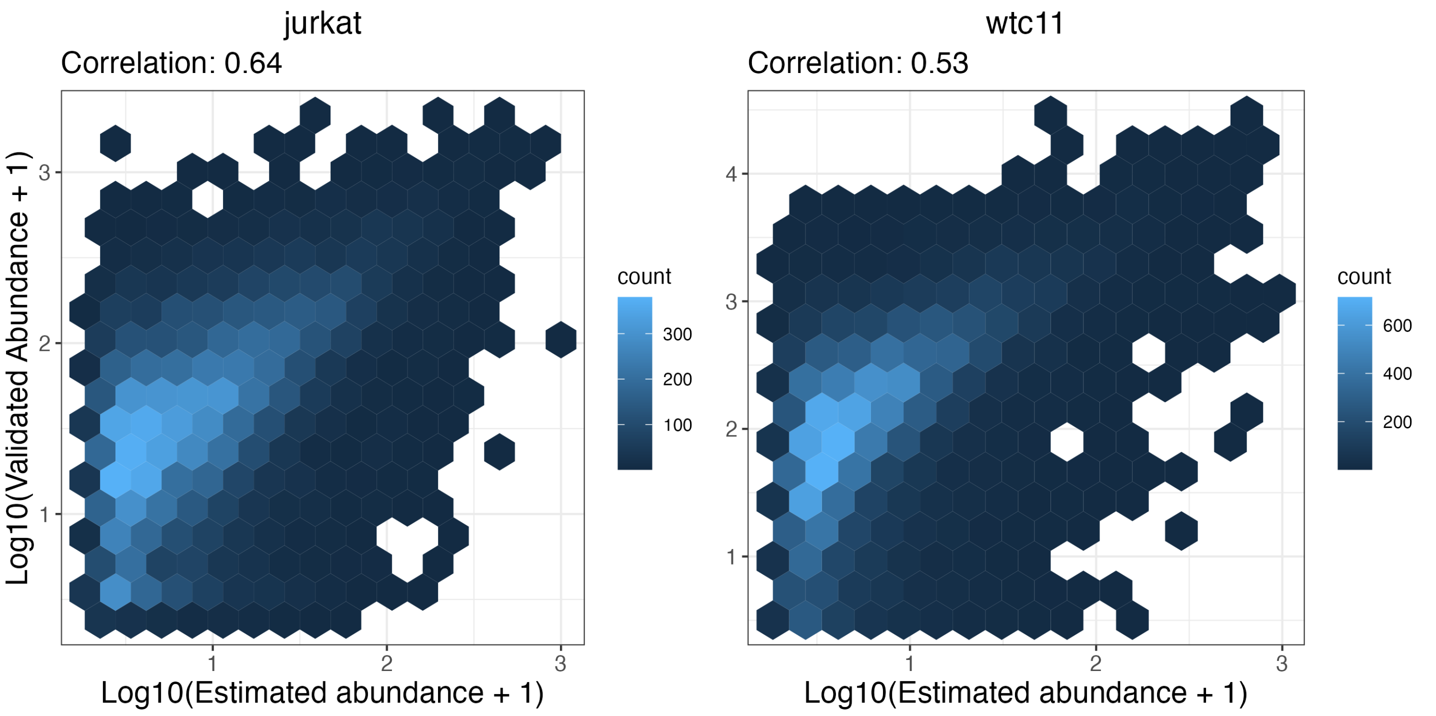


**Supplementary Figure 1:** Hexbin plot for the log10 protein isoform abundances (i.e., log10(abundance + 1)), estimated from *IsoBayes* (x axis), and found in the validation set (y axis). In each cell line, we considered results from all protesease. Left: *jurkat* dataset; right: *WTC-11* dataset.

###### 3.1.2 Gene-level results

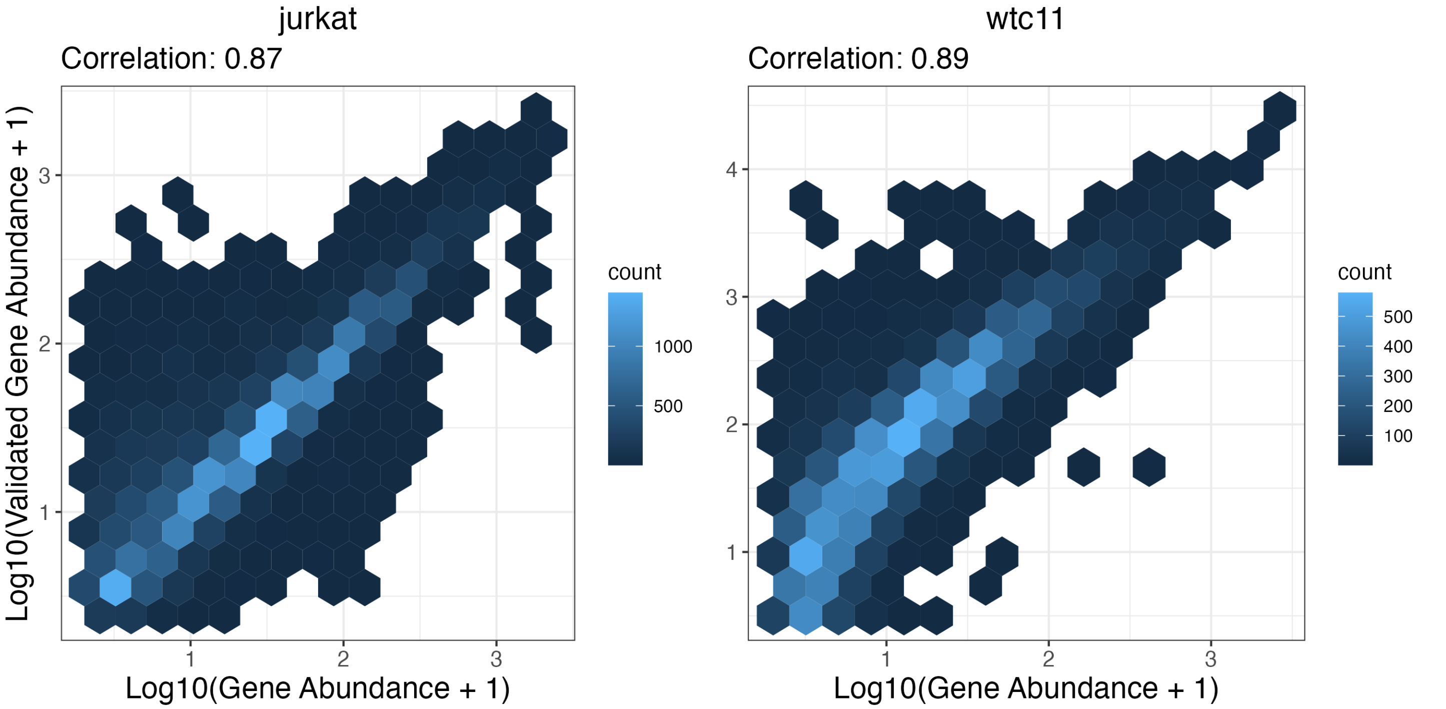


**Supplementary Figure 2:** Hexbin plot for the log10 protein gene abundances (i.e., log10(abundance + 1)), estimated from *IsoBayes_mRNA* (x axis), and found in the validation set (y axis). In each cell line, we considered results from all protesease. Left: *jurkat* dataset; right: *WTC-11* dataset.

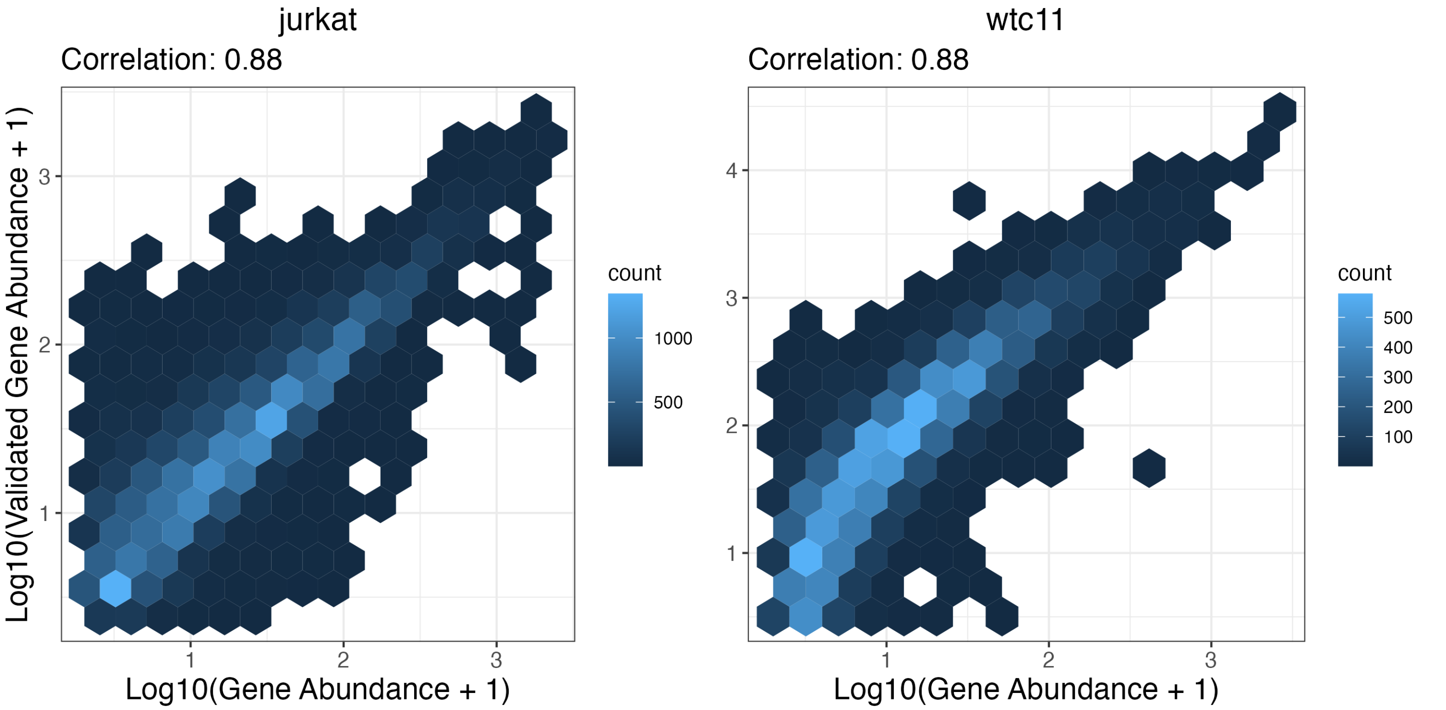


**Supplementary Figure 3:** Hexbin plot for the log10 protein gene abundances (i.e., log10(abundance + 1)), estimated from *IsoBayes* (x axis), and found in the validation set (y axis). In each cell line, we considered results from all protesease. Left: *jurkat* dataset; right: *WTC-11* dataset.

##### 3.1.3 log2-FCs

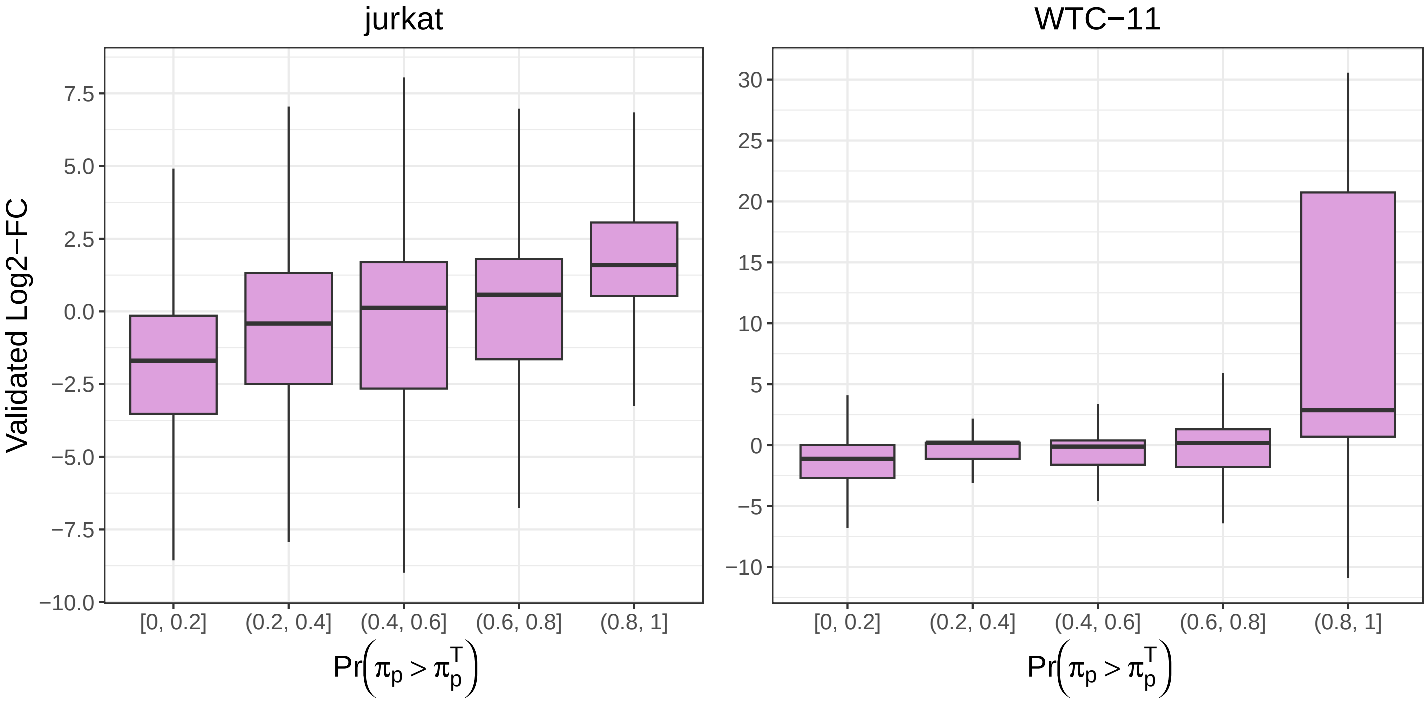


**Supplementary Figure 4:** Boxplot of the stabilized log2-FCs between protein and transcript relative abundances, identified in the validated set, stratified based on the probability, estimated by *IsoBayes_mRNA*, that isoform relative abundances are higher at the protein- than at then transcript-level. In each cell line, we considered results from all protesease. Left: *jurkat* dataset; right: *WTC-11* dataset.

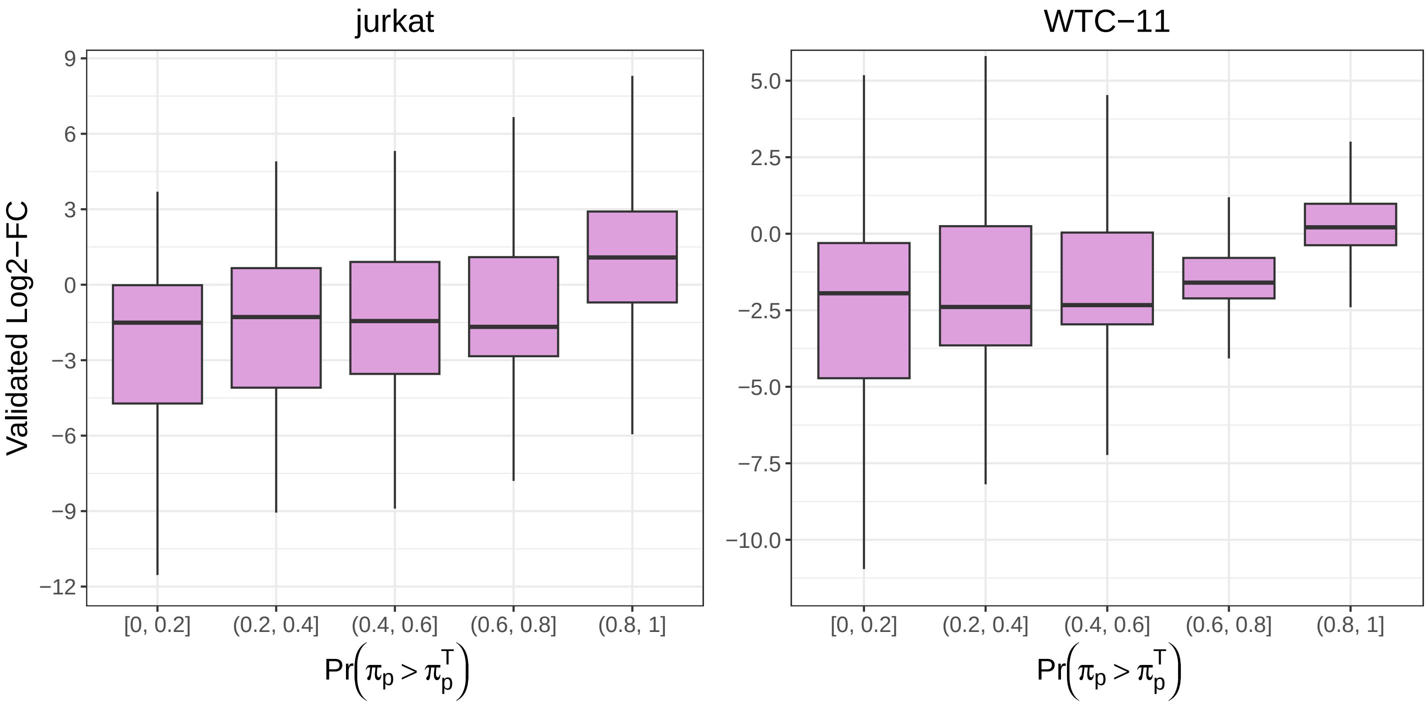


**Supplementary Figure 5:** Boxplot of the stabilized log2-FCs between protein and transcript relative abundances, identified in the validated set, stratified based on the probability, estimated by *IsoBayes*, that isoform relative abundances are higher at the protein- than at then transcript-level. In each cell line, we considered results from all protesease. Left: *jurkat* dataset; right: *WTC-11* dataset.

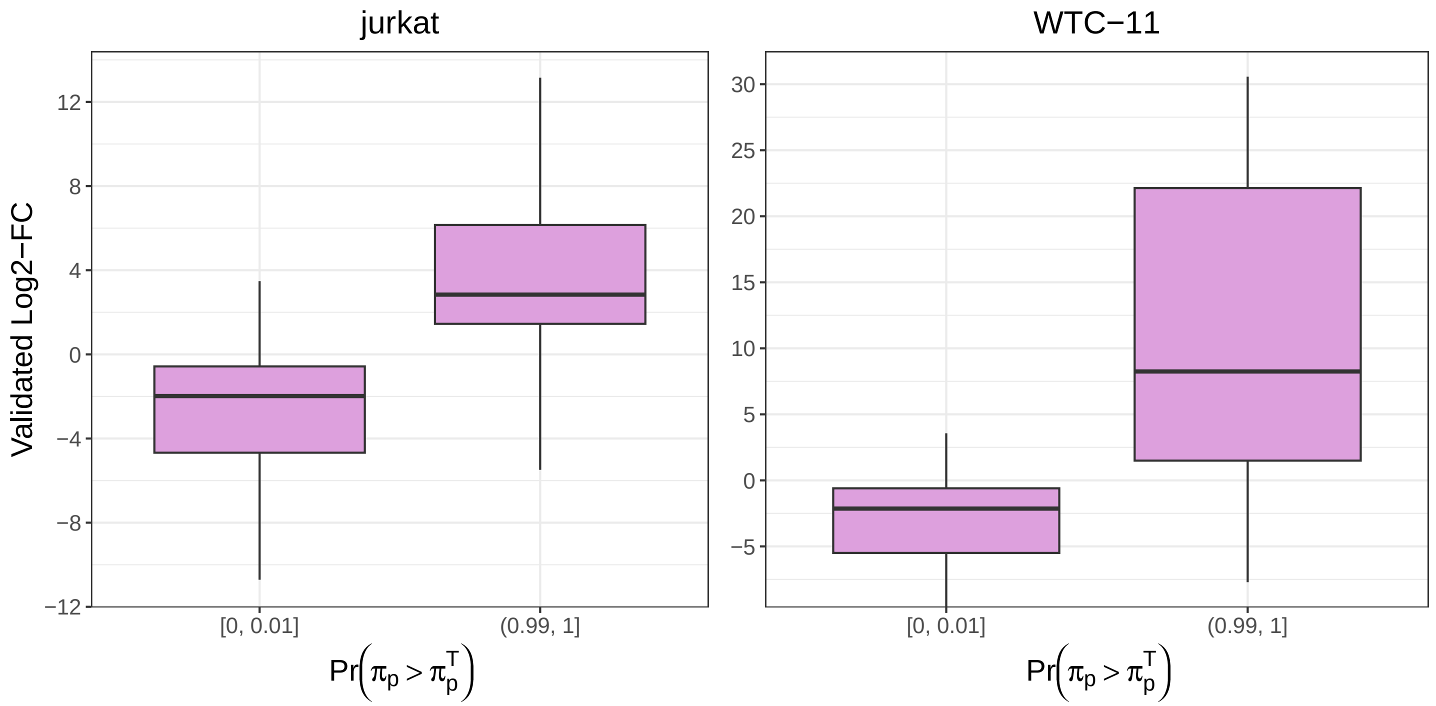


**Supplementary Figure 6:** Boxplot of the stabilized log2-FCs between protein and transcript relative abundances, identified in the validated set, stratified based on the probability, estimated by *IsoBayes*, that isoform relative abundances are higher at the protein- than at then transcript-level. Small estimated probabilities (below 0.01) are mainly associated to negative log2-FCs in the validation set; conversely, large estimated probabilities (above 0.99) typically lead to positive log2-FCs in the validation set. In each cell line, we considered results from all protesease. Left: *jurkat* dataset; right: *WTC-11* dataset.

##### 3.1.4 Memory usage

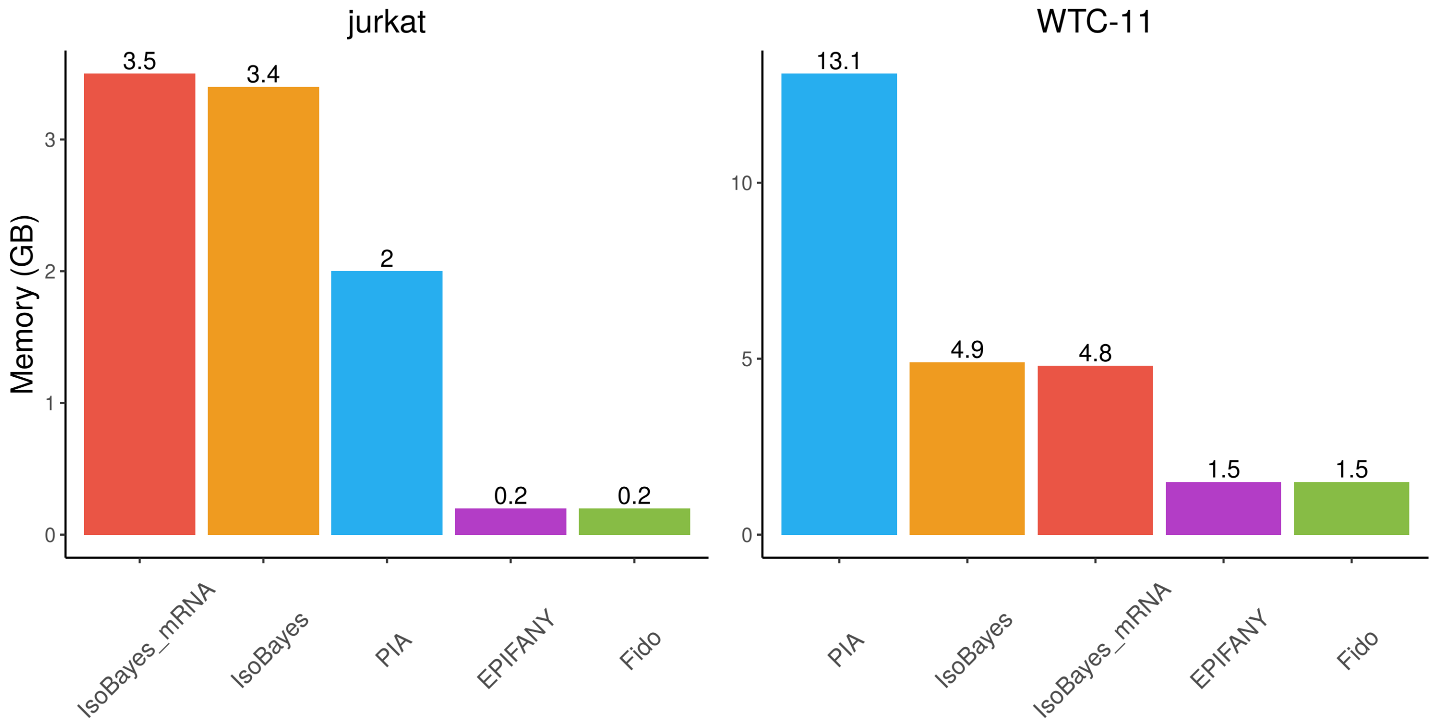


**Supplementary Figure 7:** Average (across proteases) of the maximum memory required by each method, expressed in gigabytes (GB). Left: *jurkat* dataset; right: *WTC-11* dataset.

#### 3.2 Isoforms without unique peptides

##### 3.2.1 Isoform-level results

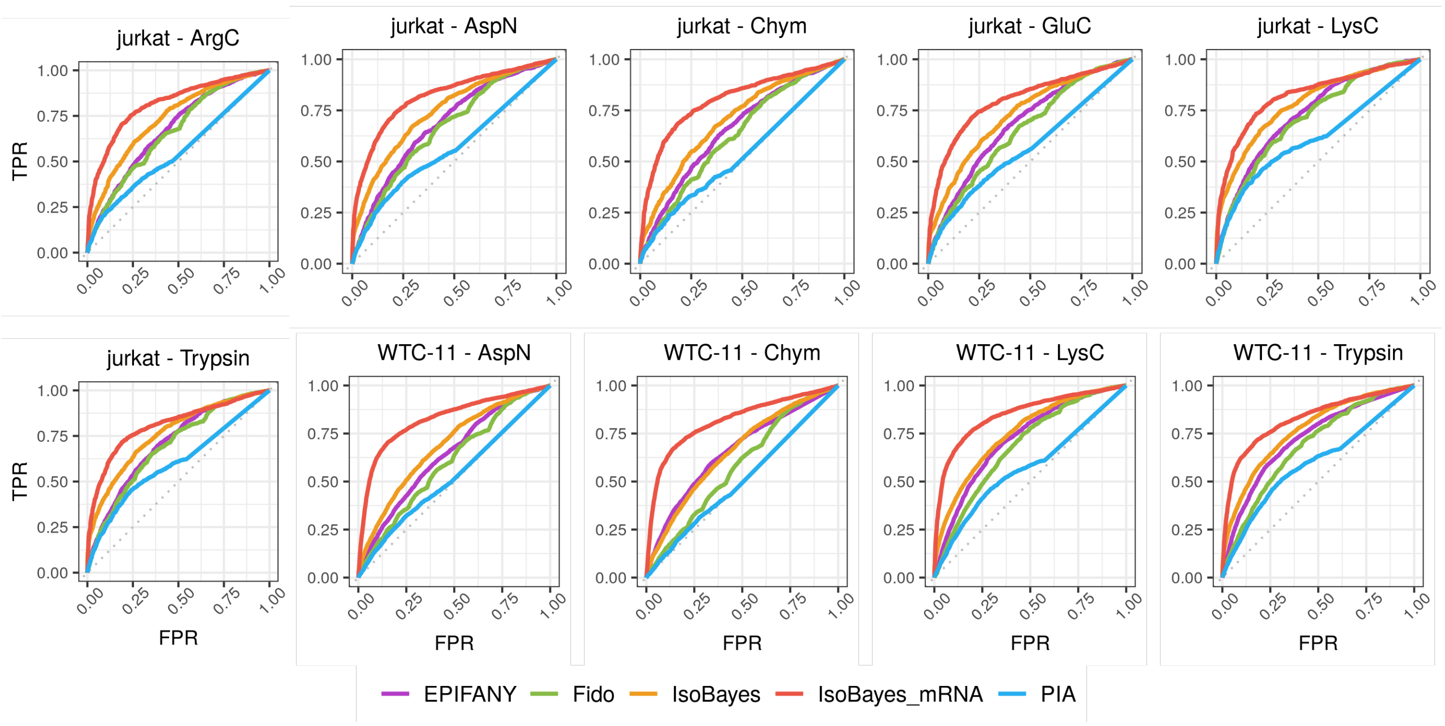


**Supplementary Figure 8:** Receiver operating characteristic (ROC) curves for the detection of protein isoforms in each real dataset, computed on the subset of protein isoforms solely associated to shared peptides.

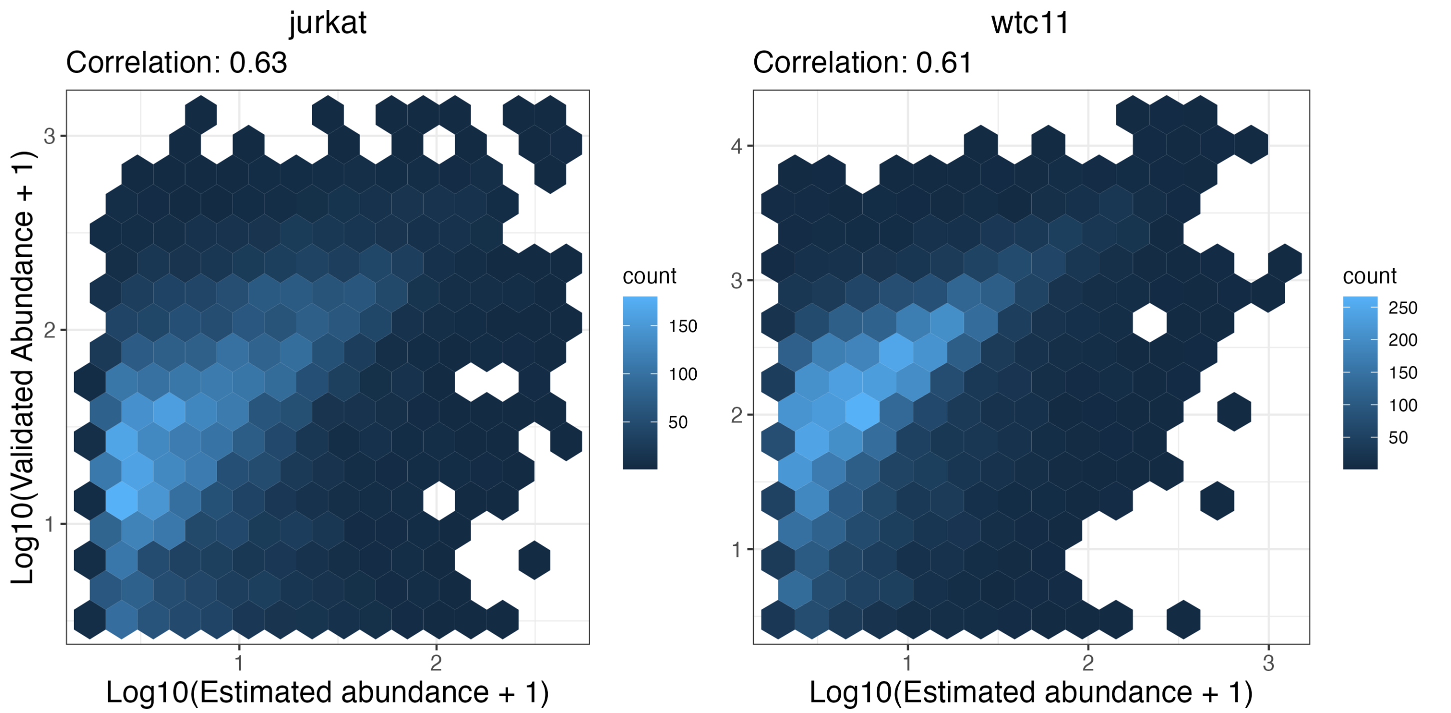


**Supplementary Figure 9:** Hexbin plot for the log10 protein isoform abundances (i.e., log10(abundance + 1)), estimated from *IsoBayes_ mRNA* (x axis), and found in the validation set (y axis), computed on the subset of protein isoforms solely associated to shared peptides. In each cell line, we considered results from all protesease. Left: *jurkat* dataset; right: *WTC-11* dataset.

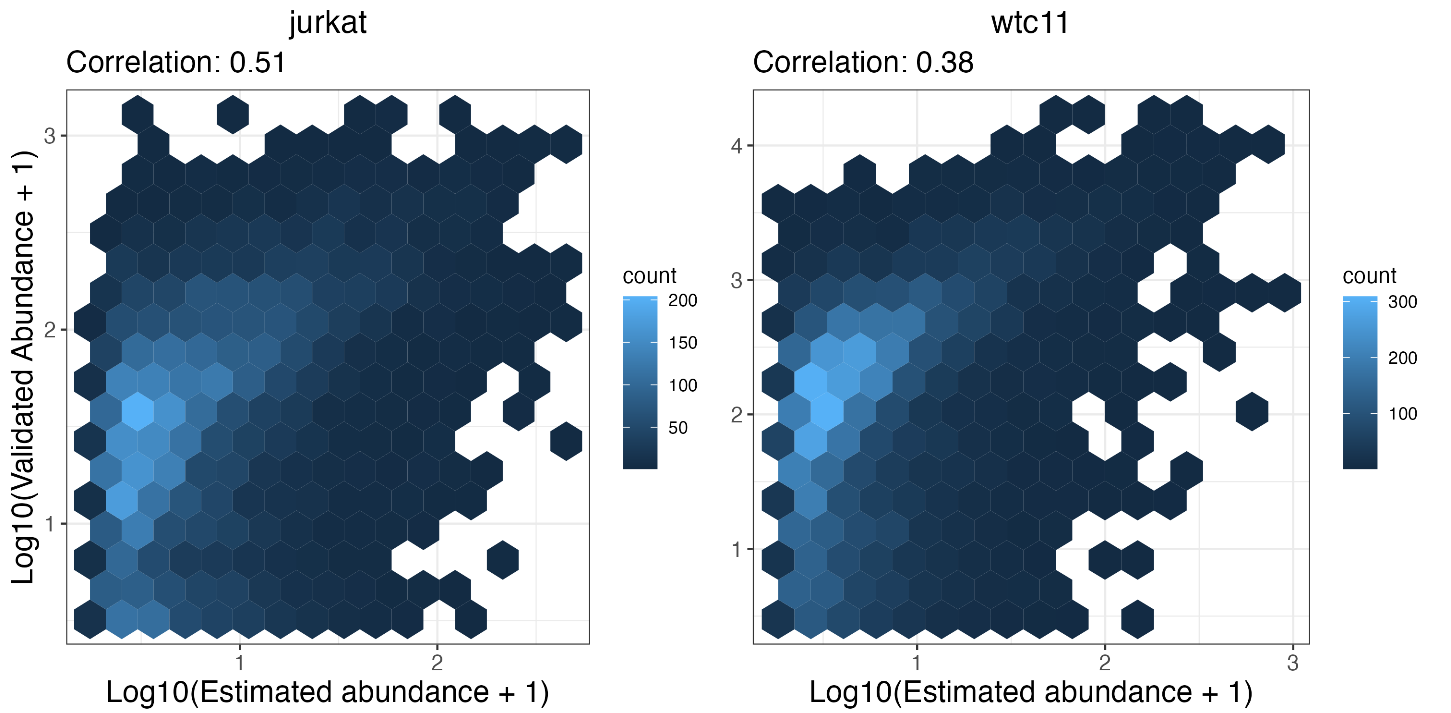


**Supplementary Figure 10:** Hexbin plot for the log10 protein isoform abundances (i.e., log10(abundance + 1)), estimated from *IsoBayes* (x axis), and found in the validation set (y axis), computed on the subset of protein isoforms solely associated to shared peptides. In each cell line, we considered results from all protesease. Left: *jurkat* dataset; right: *WTC-11* dataset.

##### 3.2.2 log2-FCs

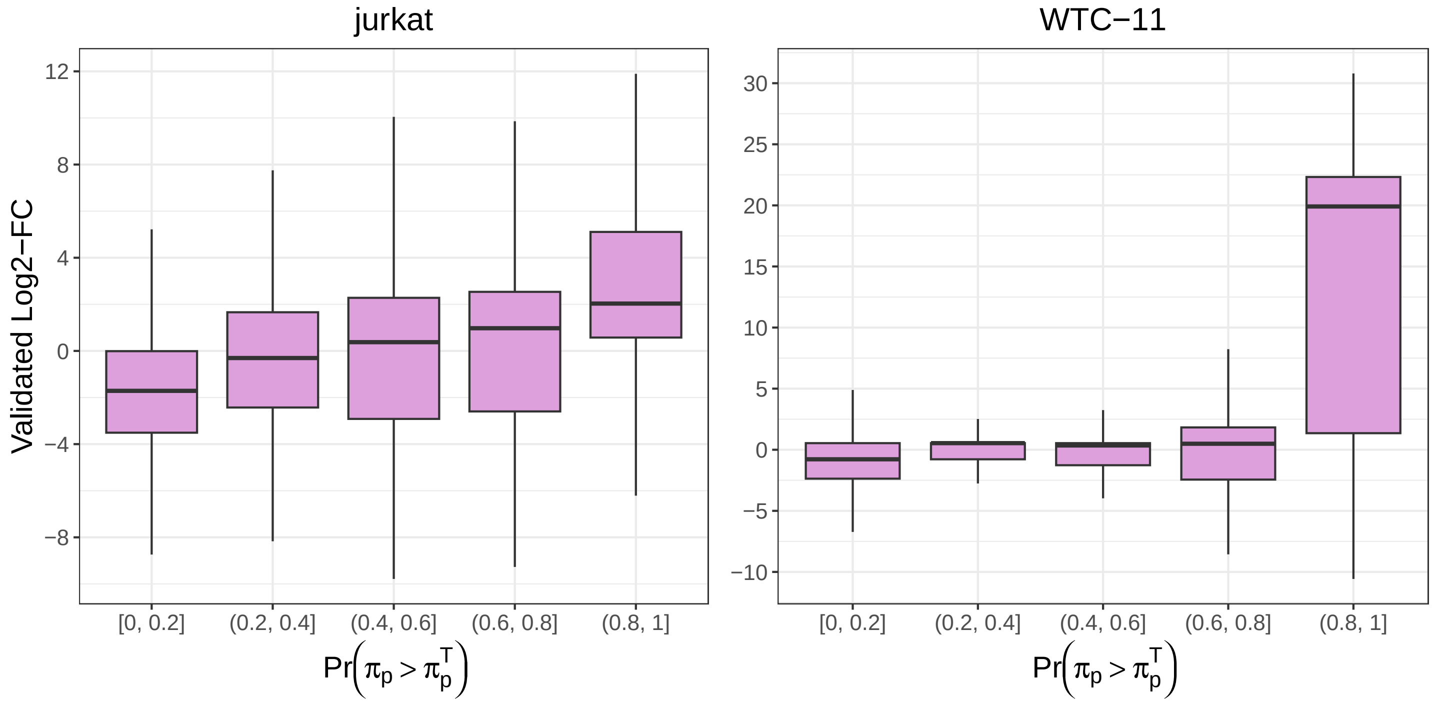


**Supplementary Figure 11:** Boxplot of the stabilized log2-FCs between protein and transcript relative abundances, identified in the validated set, stratified based on the probability, estimated by *IsoBayes_mRNA*, that isoform relative abundances are higher at the protein- than at then transcript-level, computed on the subset of protein isoforms solely associated to shared peptides. In each cell line, we considered results from all protesease. Left: *jurkat* dataset; right: *WTC-11* dataset.

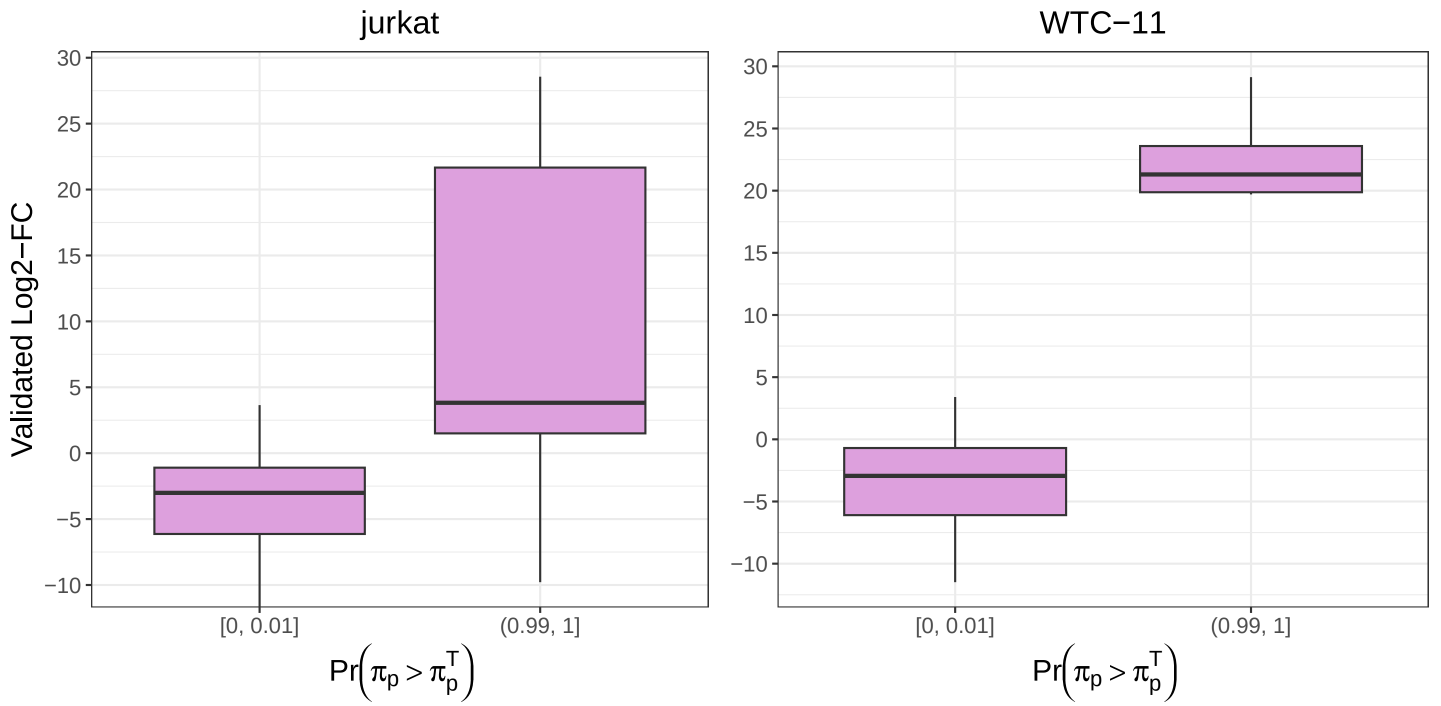


**Supplementary Figure 12:** Boxplot of the stabilized log2-FCs between protein and transcript relative abundances, identified in the validated set, stratified based on the probability, estimated by *IsoBayes_mRNA*, that isoform relative abundances are higher at the protein- than at then transcript-level, computed on the subset of protein isoforms solely associated to shared peptides. Small estimated probabilities (below 0.01) are mainly associated to negative log2-FCs in the validation set; conversely, large estimated probabilities (above 0.99) typically lead to positive log2-FCs in the validation set. In each cell line, we considered results from all protesease. Left: *jurkat* dataset; right: *WTC-11* dataset.

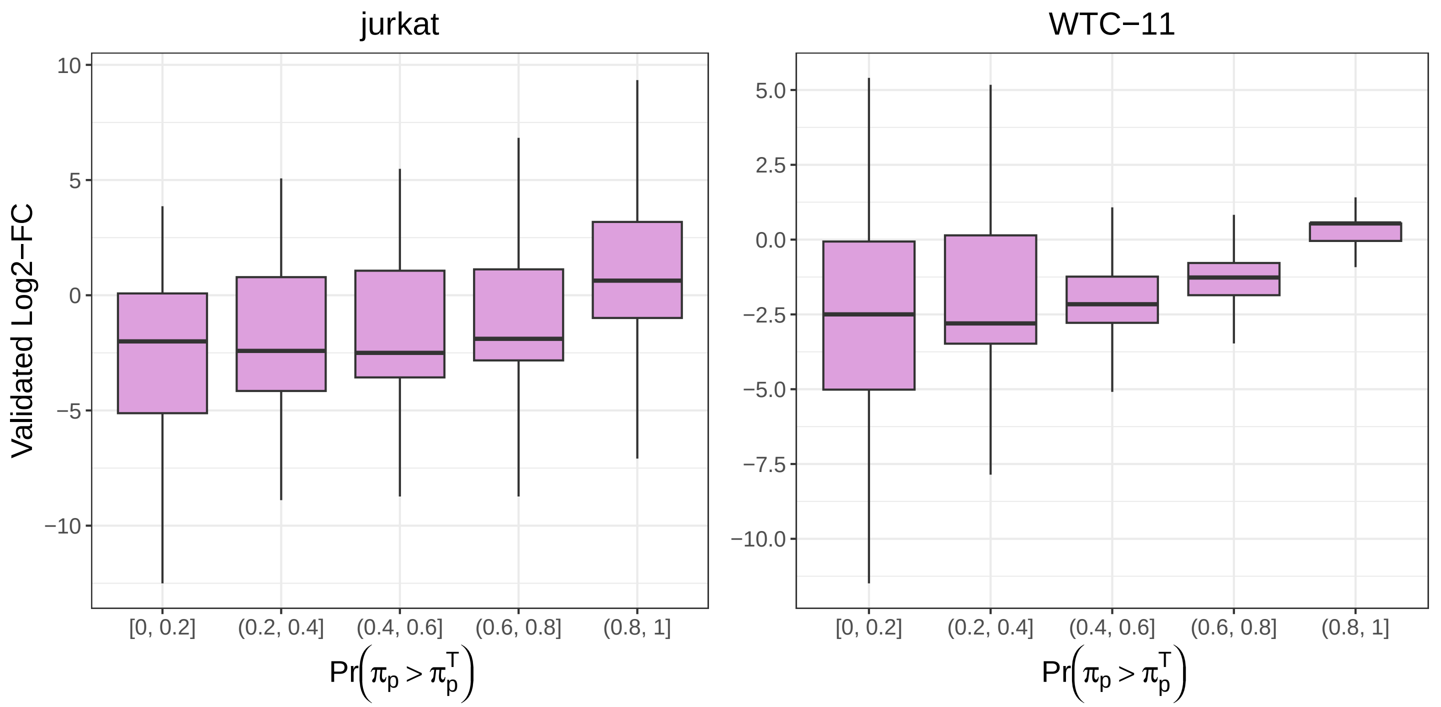


**Supplementary Figure 13:** Boxplot of the stabilized log2-FCs between protein and transcript relative abundances, identified in the validated set, stratified based on the probability, estimated by *IsoBayes*, that isoform relative abundances are higher at the protein- than at then transcript-level, computed on the subset of protein isoforms solely associated to shared peptides. In each cell line, we considered results from all protesease. Left: *jurkat* dataset; right: *WTC-11* dataset.

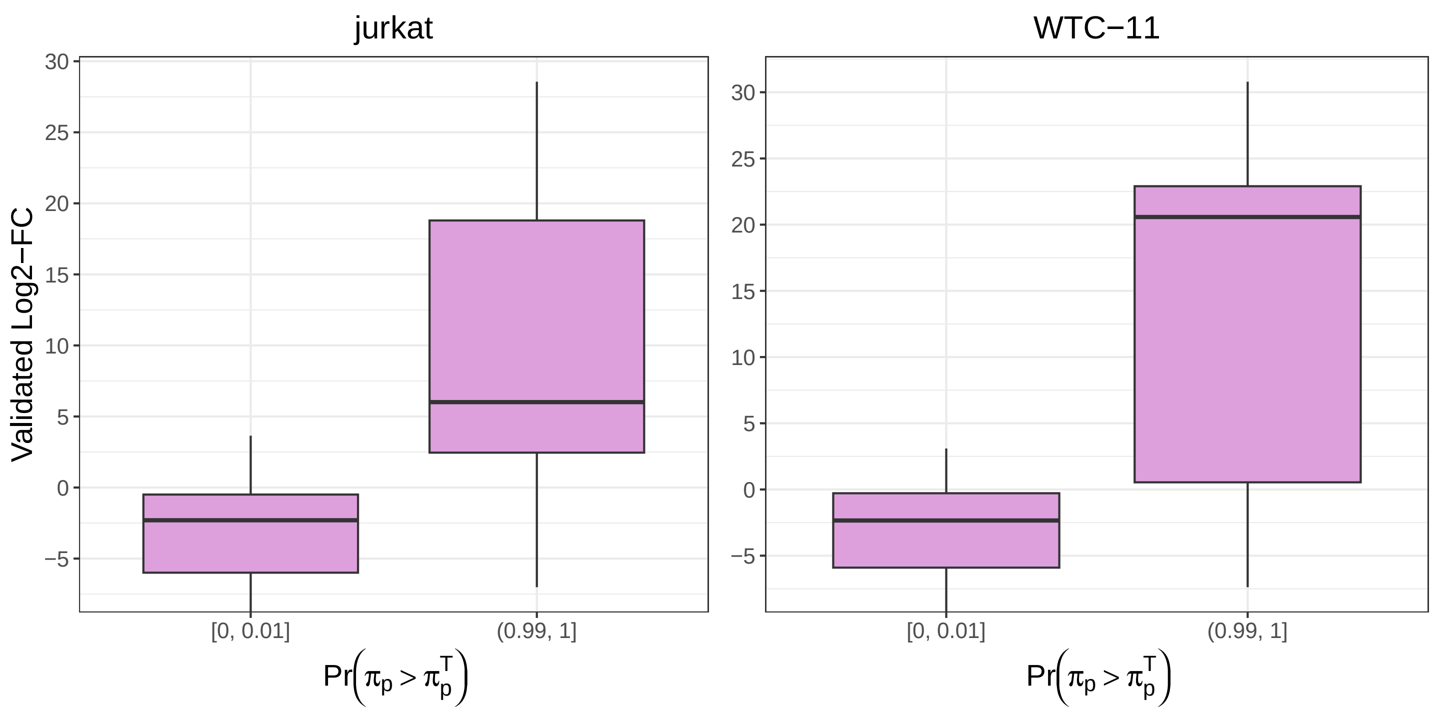


**Supplementary Figure 14:** Boxplot of the stabilized log2-FCs between protein and transcript relative abundances, identified in the validated set, stratified based on the probability, estimated by *IsoBayes*, that isoform relative abundances are higher at the protein- than at then transcript-level, computed on the subset of protein isoforms solely associated to shared peptides. Small estimated probabilities (below 0.01) are mainly associated to negative log2-FCs in the validation set; conversely, large estimated probabilities (above 0.99) typically lead to positive log2-FCs in the validation set. In each cell line, we considered results from all protesease. Left: *jurkat* dataset; right: *WTC-11* dataset.

#### 3.3 Robustness to input data

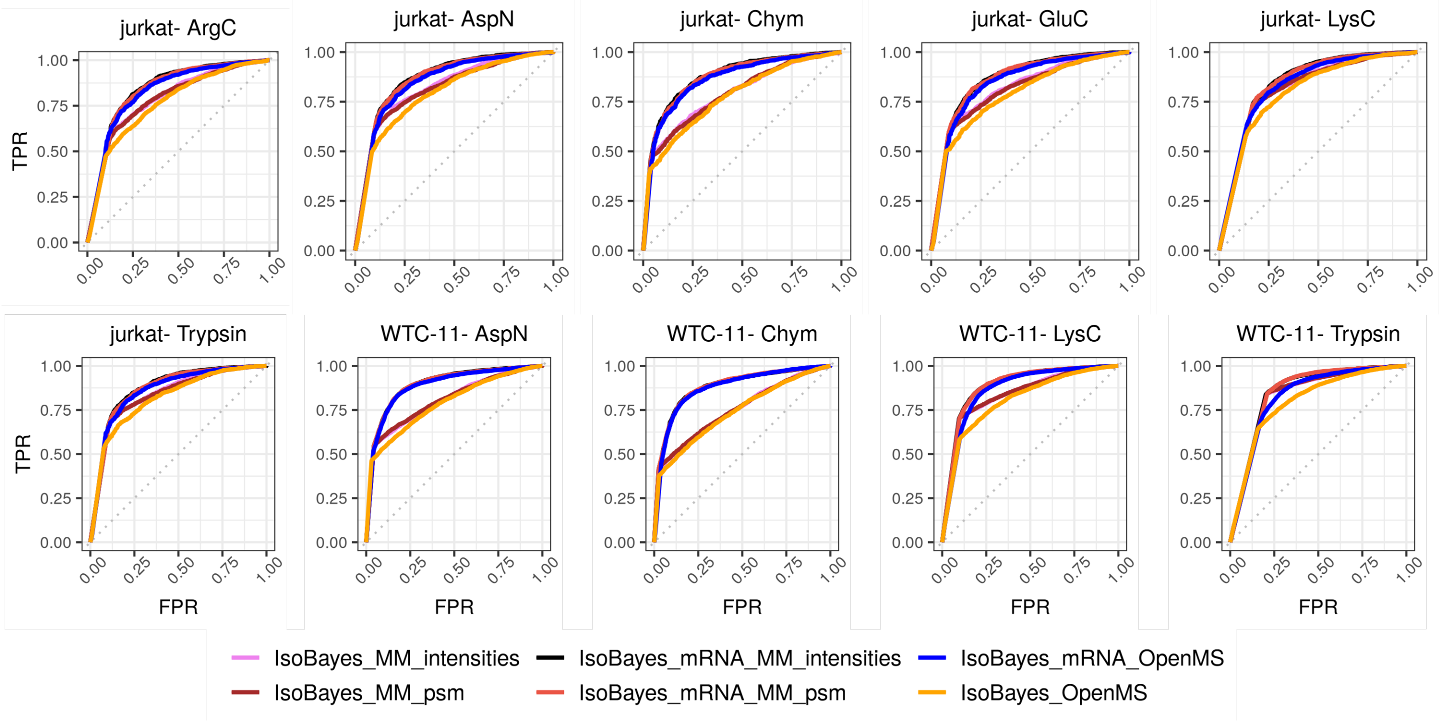


**Supplementary Figure 15:** Receiver operating characteristic (ROC) curves for the detection of protein isoforms in each real dataset, for *IsoBayes* and *IsoBayes_mRNA*, computed on three input data: i) PSM counts from *OpenMS' Percolator*; ii) PSM counts from *MetaMorpheus*; iii) peptide intensities from *MetaMorpheus*. Since slightly different peptides are detected between *MetaMorpheus* and *Percolator*, for a fair comparison, here we removed the minority of isoforms which can be analyzed with one tool only, and not with the other one.

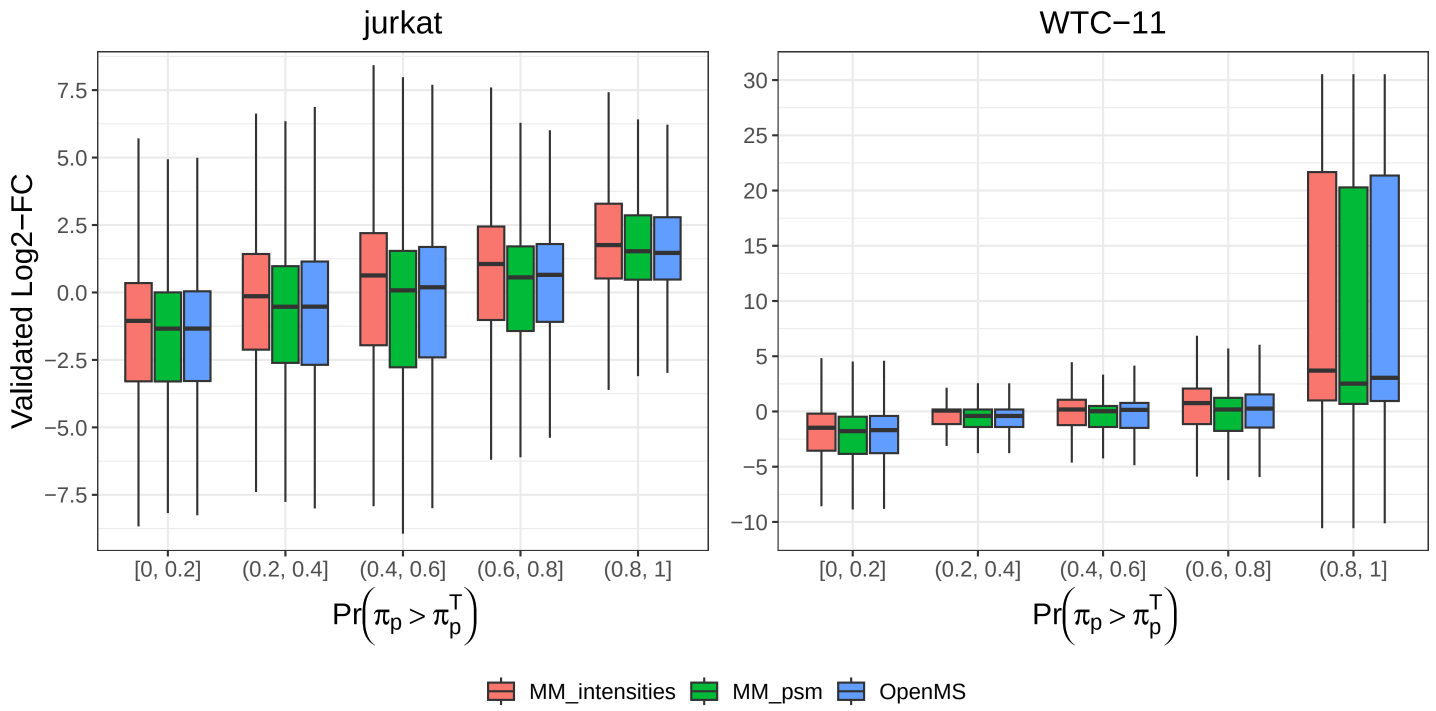


**Supplementary Figure 16:** Boxplot of the stabilized log2-FCs between protein and transcript relative abundances, identified in the validated set, stratified based on the probability, estimated by *IsoBayes_mRNA*, that isoform relative abundances are higher at the protein- than at then transcript-level. The three colours refer to the three inputs: i) PSM counts from *OpenMS*' *Percolator* ("OpenMS"); ii) PSM counts from *MetaMorpheus* ("MM_PSM"); iii) peptide intensities from *MetaMorpheus*("MM_intensities"). In each cell line, we considered results from all protesease. Left: *jurkat* dataset; right: *WTC-11* dataset. Since slightly different peptides are detected between *MetaMorpheus* and *Percolator*, for a fair comparison, here we removed the minority of isoforms which can be analyzed with one tool only, and not with the other one.

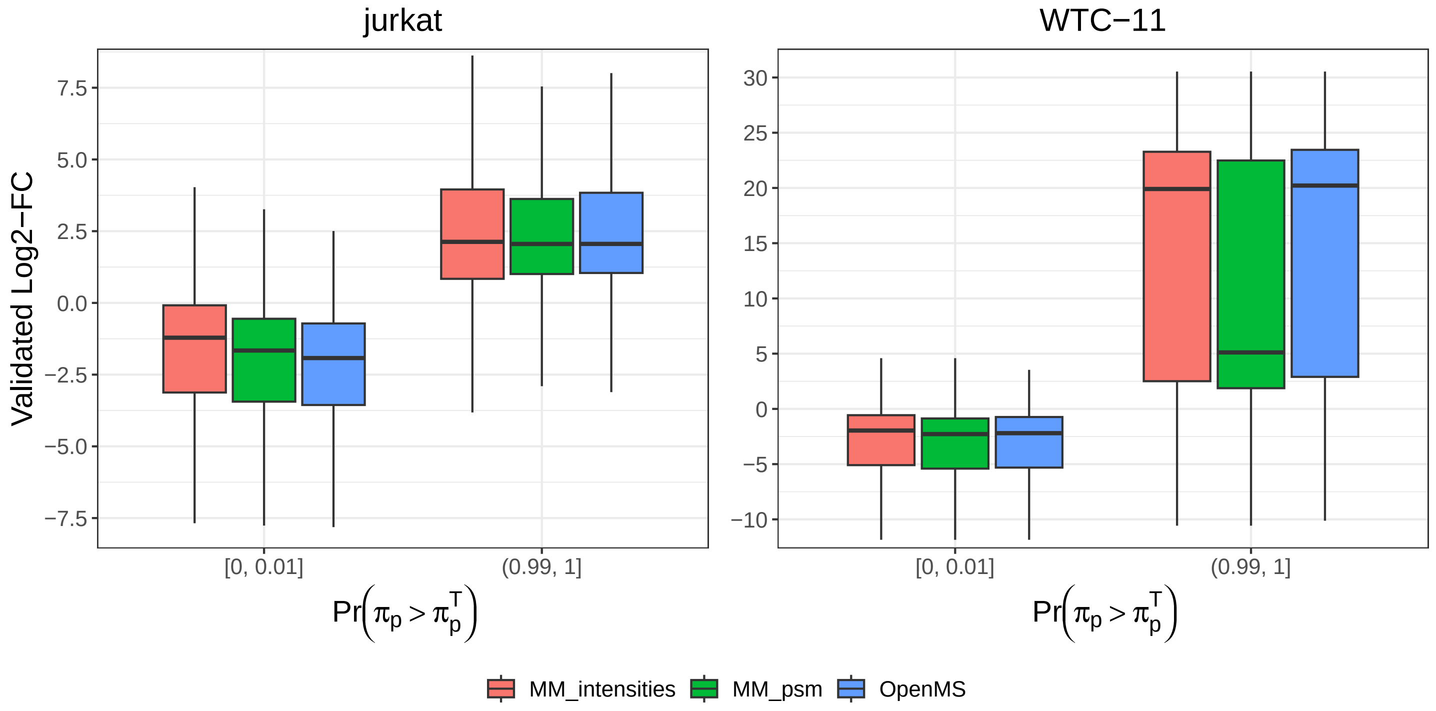


**Supplementary Figure 17:** Boxplot of the stabilized log2-FCs between protein and transcript relative abundances, identified in the validated set, stratified based on the probability, estimated by *IsoBayes_mRNA*, that isoform relative abundances are higher at the protein- than at then transcript-level. The three colours refer to the three inputs: i) PSM counts from *OpenMS*' *Percolator* ("OpenMS"); ii) PSM counts from *MetaMorpheus* ("MM_PSM"); iii) peptide intensities from *MetaMorpheus*("MM_intensities"). In each cell line, we considered results from all protesease. Left: *jurkat* dataset; right: *WTC-11* dataset. Since slightly different peptides are detected between *MetaMorpheus* and *Percolator*, for a fair comparison, here we removed the minority of isoforms which can be analyzed with one tool only, and not with the other one.

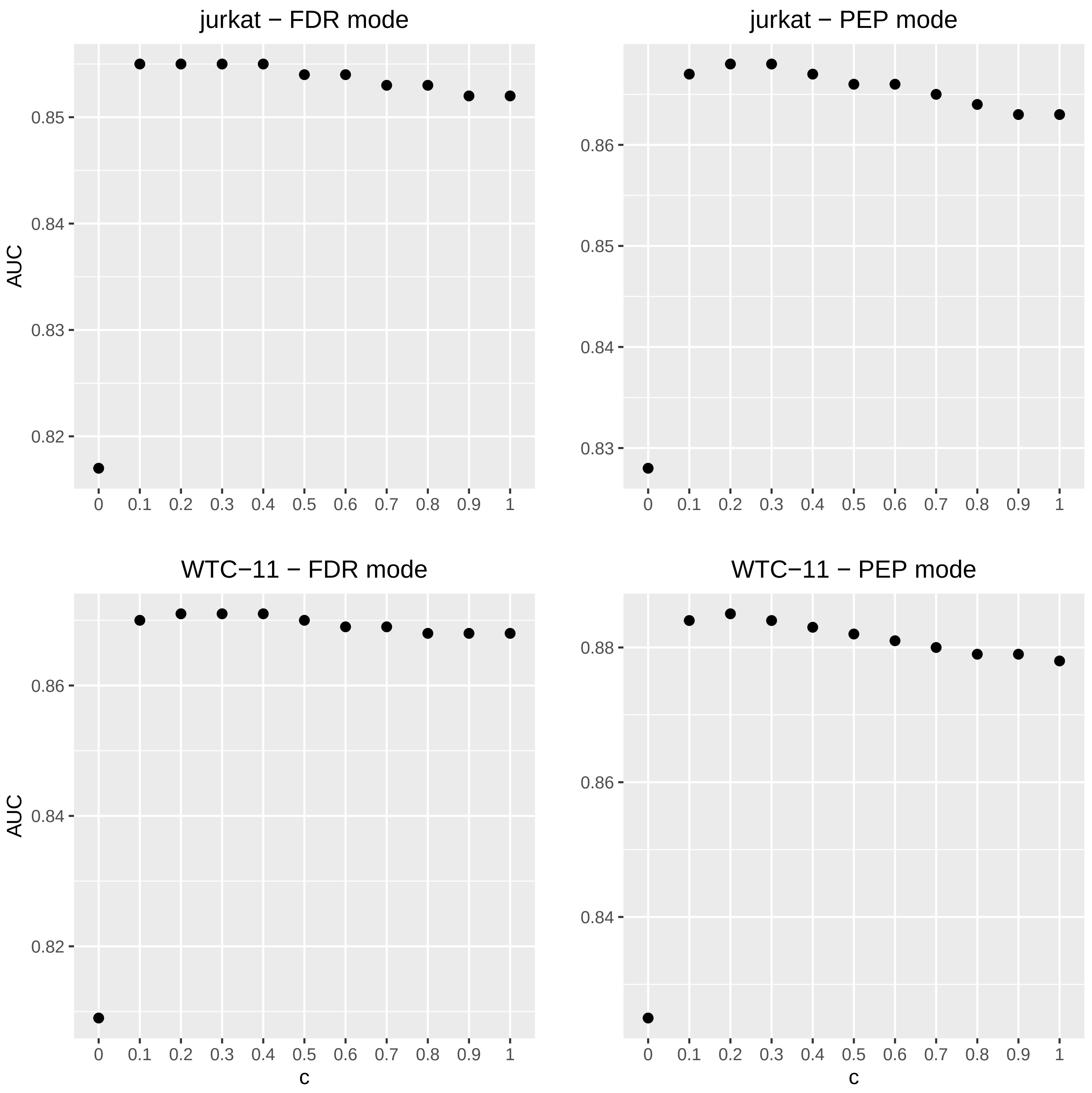


**Supplementary Figure 18:** Area under the curve (AUC) depending on the strength of the informative prior (mRNA relative abundances). On the x-axis, 0 indicates that mRNA was not used to formulate the informative prior (i.e., $\delta_1 = \cdots = \delta_P = 1$), while 1 denotes that mRNA and protein data have the same weight in the posterior distribution of $\pi|X, \delta$. By default, in *IsoBayes* "c" is set to 0.1, which in general provides the highest AUC in our benchmarks, as visible in the image. Top row: *jurkat* data; bottom row: *WTC-11* data. Left panels: PEP mode; right panels: FDR mode. AUC values represent averages across the AUC values obtained on the six *jurkat* and four *WTC-11* proteases.

### References


[1] S. K. Solntsev, M. R. Shortreed, B. L. Frey, and L. M. Smith. Enhanced global post-translational modification discovery with MetaMorpheus. *Journal of proteome research*, 17(5):1844–1851, 2018.

[2] J. R. Wiśniewski. Filter-aided sample preparation for proteome analysis. *Microbial proteomics: methods and protocols*, pages 3–10, 2018.
